# A tonically active master neuron modulates mutually exclusive motor states at two timescales

**DOI:** 10.1101/2022.04.06.487231

**Authors:** Jun Meng, Tosif Ahamed, Bin Yu, Wesley Hung, Sonia EI Mouridi, Zezhen Wang, Yongning Zhang, Quan Wen, Thomas Boulin, Shangbang Gao, Mei Zhen

## Abstract

Continuity of behaviors requires animals to make smooth transitions between successive and mutually exclusive behavioral states. Neural principles that govern these transitions are not well understood. *C. elegans* spontaneously switch between two opposite motor states, forward and backward movement, a phenomenon long thought to reflect the reciprocal inhibition between two interneurons that separately gate the forward and backward motor circuits, AVB and AVA. Combining experimental data and mathematical modeling, we report that spontaneous forward and backward locomotion and their corresponding motor circuits are not separately controlled. AVA and AVB are neither functionally equivalent nor strictly reciprocally inhibitory. Instead, while AVA phasically inhibits the forward promoting interneuron AVB at a fast timescale, it maintains a tonic, extrasynaptic excitation on AVB over the longer timescale. AVA’s depolarized spontaneous membrane potential is necessary for this tonic excitation. We propose a new, master neuron model for locomotion. AVA, with tonic and phasic activity of opposite polarities on different time scales, acts as a master neuron to break the symmetry between the underlying forward and backward motor circuits. This offers a parsimonious solution for sustained locomotion consisted of mutually exclusive motor states.

**Teaser:** A tonically active *C. elegans* premotor interneuron functions as the master neuron that underlies continuous modulation of forward and backward movement to ensure smooth transitions between the two opposing motor states.

## Introduction

Animal behaviors are displayed as a sequence of discrete, often mutually-exclusive motor states (*1*–*11*), such as forward and backward locomotion. Prevailing models for selecting and switching between mutually-exclusive circuit activity patterns include the winner-take-all and reciprocal inhibitory networks (*12*–*16*). However, at shorter timescales, gradual transitions observed between successive motor states may also require continuous, albeit rapid modulation of dynamical variables. For example, locomotion velocity undergoes a continuous change in magnitude, not just a sudden change in sign, when switching between forward and backward movement (*17*–*21*). Limb movement is correlated with rotational, instead of alternating spinal neural network activity dynamics(*22*). However, circuit solutions for continuous modulation that underlie smooth behavioral state transitions are not well understood.

*C. elegans* exhibits two mutually exclusive motor states: forward movement and backward movement, propelled by the posteriorly and anteriorly propagating undulatory bending waves, respectively (*23*). In isotropic environments, they spontaneously switch between the forward and backward motor states (*23*). The long-standing model for selecting and switching between these motor states involves a reciprocally inhibitory circuit motif between two separate modules (*5*) (Fig. 1A).

**Figure 1.**
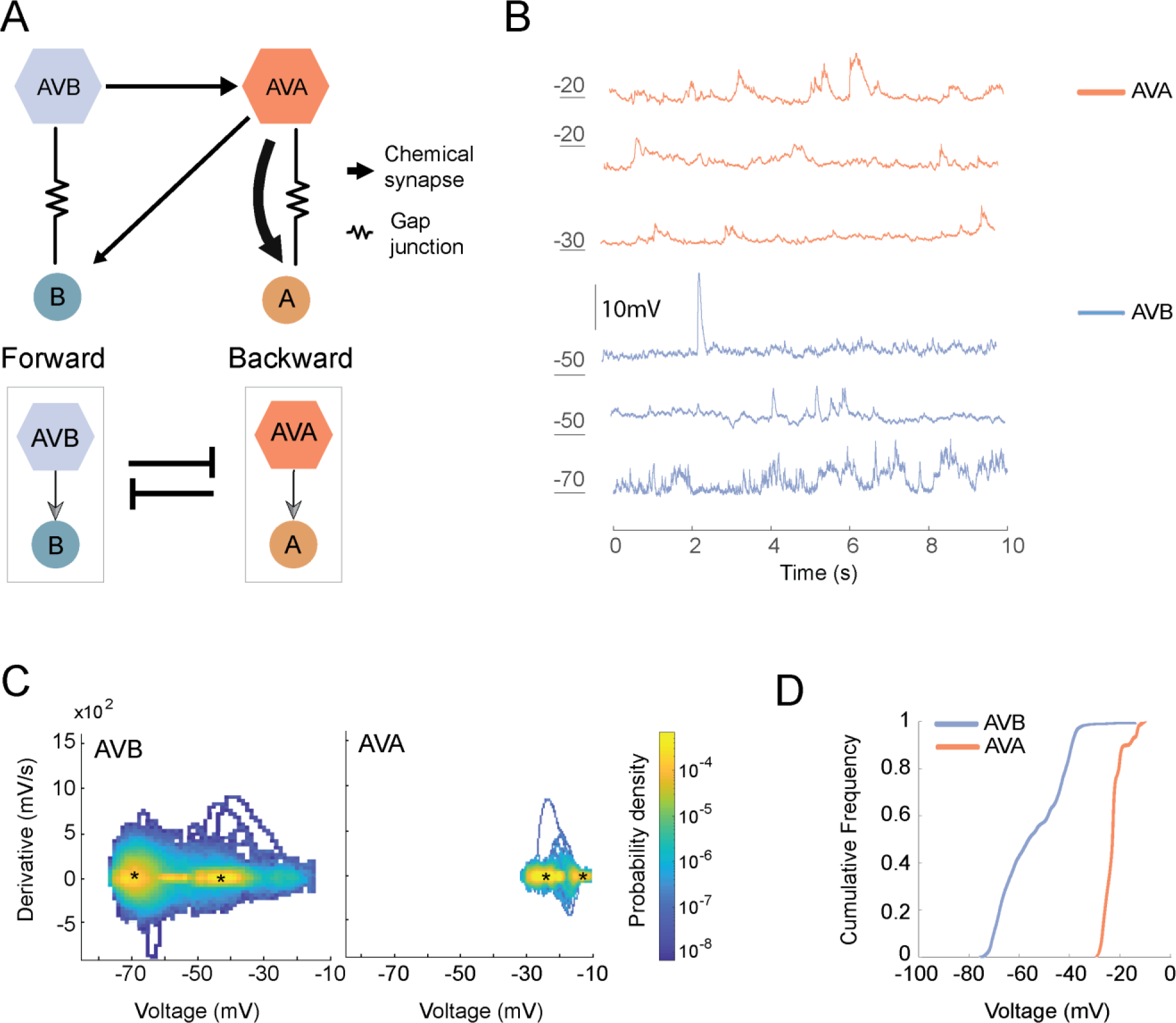
AVA and AVB have distinct membrane potential ranges and dynamics. **A.** (Upper) Schematics of the adult anatomic wiring diagram for premotor interneurons and motor neurons implicated in directional movement, adapted from(*24*, *59*, *68*); (Lower) Schematics of previously proposed functional relationship of these neurons(*27*– *31*). **B.** Example traces of spontaneous membrane potential of AVA and AVB neurons by *in situ* whole-cell intracellular recording configuration. **C.** The dynamic range of spontaneous membrane potential for AVB and AVA, is represented by the distribution of spontaneous membrane voltage and its derivatives. * denotes regimes of membrane potentials that might correlate with bi-stability described in previous studies(*38*, *43*, *44*). AVB potential drifts over a wider range than AVA (X-axis) and exhibits more frequent but smaller transients than AVA (Y-axis). **D.** Cumulative curves of the distribution of spontaneous membrane potential of AVB and AVA. AVA holds more depolarized membrane potential than AVB. N: 8 AVB and 7 AVA, each neuron was recorded for 60- 200 seconds.

Anatomically, two groups of non-spiking premotor interneurons, AVB and AVA, make dominant synaptic inputs onto two groups of motor neurons that separately execute the forward and backward movements (*24*–*26*) (Fig. 1A). Based on these connectivity patterns, it has been assumed that AVA and AVB also independently control backward and forward movement (*27*–*32*). Synaptic connectivity between these modules (Fig. 1A) is thought to regulate transitions and dwell times in each behavioral state (*5*). This forms the base of a phenomenological model, where the forward and backward states are treated as binary outcomes of two symmetrically wired and functionally independent circuits (*5*), and transitions between forward and backward states occur stochastically or are driven by external input (*5*).

While a reciprocal inhibition-based neuron model is intuitive, previous measurements of **A.** *C. elegans* locomotion have revealed other features not easily captured by modelling motor patterns through strictly reciprocal inhibition. Examples include the continuous decay of forward speed prior to the transition into backward movement (*18*, *23*, *33*), the period of post-stimulus arousal during which animals exhibit a higher forward velocity after an escape response (*34*, *35*), and the observation that reducing the activity of premotor interneurons of the backward module decreases both backward and forward speed (*36*).

We sought relationships between the forward and backward motor circuits when animals perform spontaneous locomotion. By examining the animal’s spontaneous motor pattern, the electrophysiological recording and calcium imaging data for AVA and AVB, we found that our experimental data can be integrated into a simple dynamical systems model. Critically, both our experimental data and the dynamical systems model reveal that AVA and AVB cannot be functionally the equivalent, independent controllers for the forward and backward motor states. Instead of being strictly reciprocally inhibitory, AVA and AVB neurons exhibit a master-slave configuration, where AVA controls AVB with opposite polarities at different timescales.

Specifically, we found that AVA inhibits AVB at short timescales while excites it at longer timescales. The former results in transient backward movement, while the latter promotes the forward motor state as a whole and transitions from backward movement to forward movement in particular. We further demonstrated that, mechanistically, the slow constitutive excitation of AVB results from the tonic acetylcholine activity of AVA due to its depolarized membrane potential.

AVA neuron has been considered to play a dedicated role in the backward motor state (*27*–*29*, *31*, *32*), commonly viewed to simply trigger a backward movement. This is likely an oversimplification due to the multi-functionality of neurons that constitute a compact nervous system. Previously, we showed that AVA both inhibits and promotes backward movement (*28*). Here, we further revealed that it functions as a master neuron for not only the backward motor states, but also forward motor states and transitions between them.

More important, conceptually, these results establish an inherent asymmetry in a motor circuit that underlies spontaneous switches between two mutually exclusive motor states. This new and parsimonious circuit solution for selecting and switching between behavioral states should have broad relevance in understanding decision-making circuits.

## Results

### Asymmetry in AVA and AVB’s membrane excitability

To begin exploring their relationships, we first examined AVA and AVB’s membrane potential by in situ electrophysiological recording (Methods) (*37*, *38*). Like most *C. elegans* neurons, AVA and AVB do not fire all-or-none action potentials. Instead, their membrane potentials fluctuate in an analog fashion (*5*, *38*–*43*). We observed that both AVA and AVB maintain its membrane potential at multiple stable states (Fig. 1B), making it non-trivial to estimate a single value for their ‘resting’ membrane potentials. For more appropriate presentation of their membrane property, we instead summarized the neuron’s voltage range by the cumulative distribution function (Fig. 1D), and applied a histogram estimate of the phase-space distribution - the joint probability distribution of the membrane voltage and its time derivative - to quantify the transient dynamics of the membrane voltage (Fig. 1C).

Our quantification of our AVA whole-cell patch clamp recording supports and expands the previous conclusion (*38*, *43*, *44*) that AVA maintains a depolarized potential at multiple stable states. Specifically, AVA’s potential ranged from approximately -30 mV to -10 mV (median -23 mV) (Fig. 1C, D). Its fluctuation exhibits two regimes with small time derivatives, one from -30 mV to -20 mV, and the other from -15 mV to -10 mV (Fig. 1C).

Our whole-cell patch clamp recording showed that AVB also exhibit multiple stable states. However, compared to AVA, AVB’s membrane potential resides at a much hyperpolarized range, from approximately -75mV to -10mV (median -57mV) (Fig. 1D). Moreover, AVB exhibits larger dynamic changes than that of AVA, at different stable states but most prominently from -75 mV to -45 mV (Fig. 1C).

The distinct membrane potential characteristics between AVA and AVB raised the possibility of their asymmetric roles in motor control. AVA, but not AVB, is likely to be tonically active: even at its baseline, AVA’s membrane potential is near the opening threshold of UNC-2 (*45*), a high voltage-gated calcium channel that promotes synaptic vesicular release (*46*–*48*).

### Endogenous manipulation of AVA’s membrane excitability

Each neuron’s distinct membrane excitability is regulated by its inward Na+ and outward K+ leak conductance (Fig. 2A). Most *C. elegans* neurons exhibit analog membrane activity that is likely fine-tuned by a large family of the K+ leak channel, K2P (Fig. 2A). To endogenously perturb AVA’s membrane potential, we genetically engineered TWK-40 (Fig. 2B), a K2P channel that we found to exhibit detectable expression from its endogenous genomic locus in three neuron classes, AVA, AVB, and DVA, with AVA expression level being the highest (Fig. 2C).

**Figure 2.**
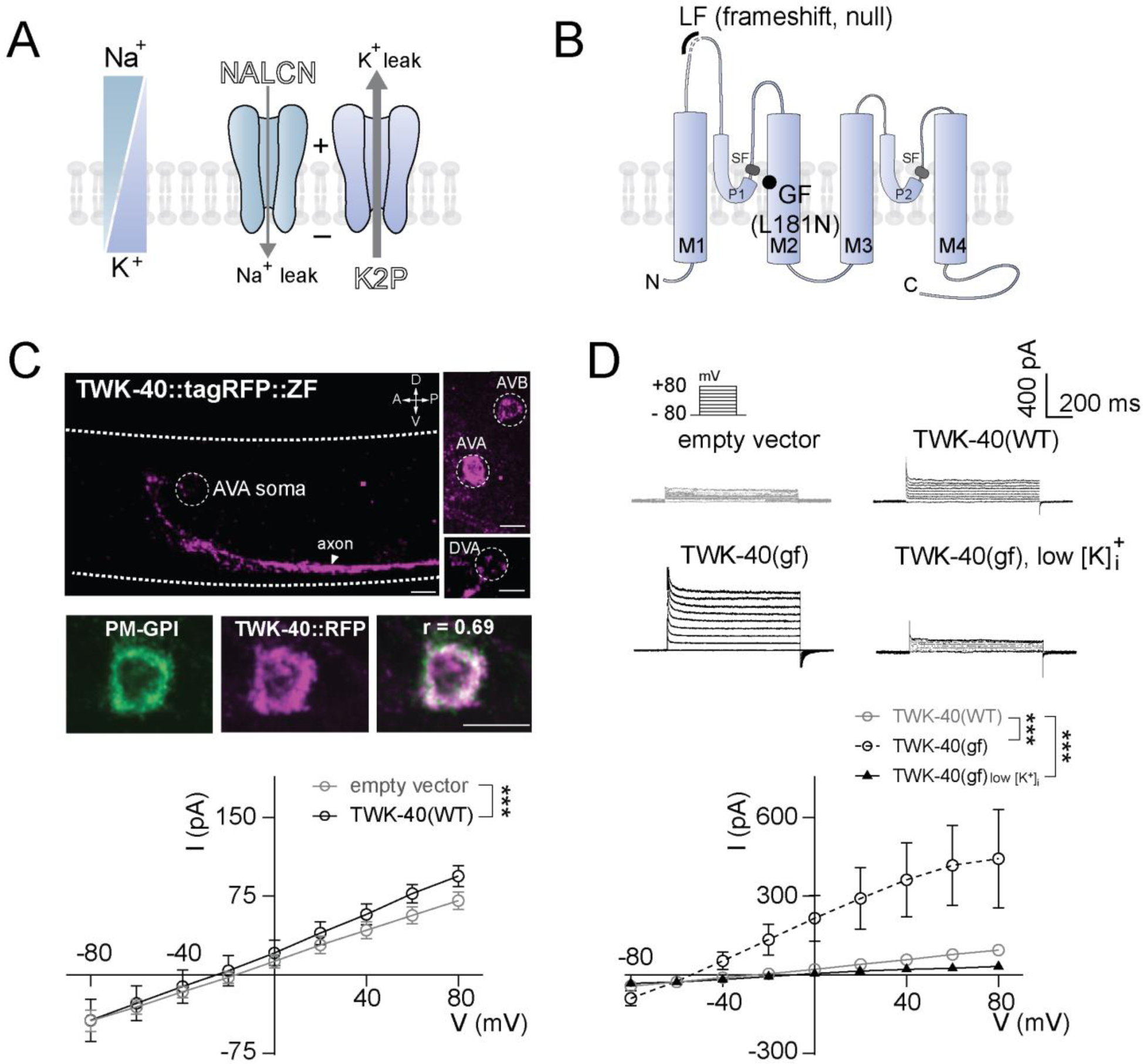
An endogenous regulator of AVA’s spontaneous membrane potential. **A.** Schematics of the inward sodium and outward potassium leak currents and respective leak channels that contribute to a cell’s membrane potential. **B.** Predicted topology of a potassium leak channel (K2P), TWK-40, denoted with genetic mutations that abolish (LF) or increase (GF) potassium leak. M: transmembrane domain; P: pore domain; SF: Selection filter; LF: loss-of-function; null: an allele (*hp834*) with a 7bp deletion that leads to truncation after the M1 domain. GF: Gain-of-function; L181N: an allele (*bln336*) with an activating mutation that promotes the potassium conductance of all K2P channels. **C.** Expression pattern of endogenously tagged TWK-40::tag RFP. (Upper) Confocal images of immunostained TWK-40::TagRFP (anti-TagRFP). Dashed lines outline the animal body; Circles denote somas. The arrowhead labels axonal processes in the ventral nerve cord. Scale bar, 2µm. (Lower) TWK-40::TagRFP (anti-TagRFP) co-localizes with a plasma membrane marker PM-GPI (anti-GFP). r: Pearson’s correlation**. D.** (Upper) Representative step currents from HEK293 cells carrying respective heterologous expression vectors by whole-cell voltage-clamp recording. (Lower) I-V curves for respective cells. Quantification showed mean currents and 95% confidence interval (CI) at +80 mV. (Left) cells expressing wild-type TWK-40 (N: 13) showed a small outward current compared to the cell with empty vector (N: 12); (Right) cells expressing TWK-40(GF) showed a large outward current (N: 10), which was diminished by reducing intracellular K+ concentration (N: 7). *: P < 0.05, **: P < 0.01, ***: P <0.001 by the Mann-Whitney U tests, with P < 0.05 considered to be statistically significant.

TWK-40 forms a functional K+ leak channel. Heterologous expression of TWK-40 in the HEK293T cells led to a small increase of the host cell’s outward leak current (Fig. 2D), similar to a few mammalian K2P channels (*49*–*51*). A gain-of-function (gf) form, which carries a mutation (L181N) that universally increases K2P channel activity (*52*, *53*), led to a large increase in the outward leak current that depends on the intracellular K+ (Fig. 1D).

We generated, then examined the effect of both loss-of-function (lf) and gain-of-function (gf) mutations in the *twk-40* genomic locus (Fig. 2B; Methods). As expected, AVA exhibited a decreased and increased outward leak current in *twk-40(lf)* and *twk-40(gf)* animals, respectively (Fig. 3A, left panel). Consistent with the effect of changing outward K+ leak, AVA’s membrane potential became further depolarized in *twk-40(lf)* mutant (∼-25mV to 0mV, median -20mV), and hyperpolarized in *twk-40(gf)* (-40mV to - 10mV, median -35mV) (Fig. 3A, right panel; Fig. 3S1). AVA-specific restoration of TWK- 40 led to increased outward leak current (Fig. 3A, left panel) and hyperpolarization (Fig. 3A, right panel; Fig. 3S1).

**Figure 3.**
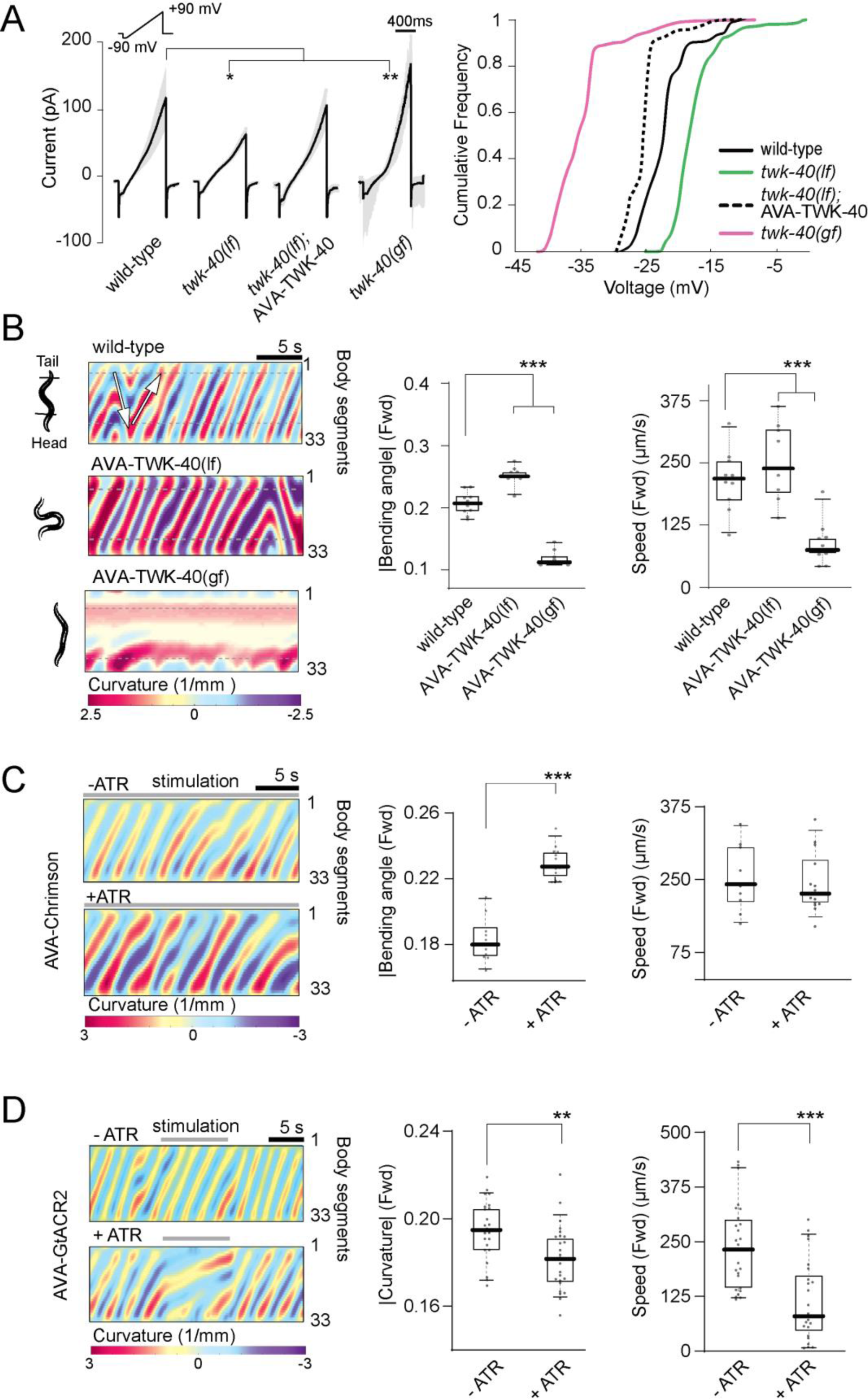
AVA’s depolarized membrane potential maintains forward movement. **A.** (Left) AVA’s leak currents recorded in whole-cell patch clamp configuration with ramping potential from -60mV to +90mV. Black traces and gray areas denote mean currents at +80mV and min/max range, respectively. N: 5 (wild-type), 8 (*twk-40(lf)*), 7 (*twk-40(lf);* AVA-TWK-40), and 7 *twk-40(gf)* cells. (Right) cumulative frequency of AVA’s spontaneous potential in respective genotypes. N: 7 (wild-type), 9 (*twk-40(lf)*), 9 (*twk-40(lf);* AVA-TWK-40) and 13 (*twk-40(gf)*) cells. Changing TWK-40’s conductance specifically in AVA led to correspondent changes in outward leak currents. **B.** (Left) Representative kymograph of bending angles of a freely moving animal of denoted genotype. Regions between dashed lines denote the animal body. Arrows illustrate directions of bending waves that correspond to either the forward (head to tail), or backward (tail to head) movements. Darker colors denote deeper bending. AVA-specific decrease (LF) or increase (GF) of TWK-40 conductance led to increased or decreased forward movement, respectively. (Right) Quantification of absolute bending angles and centroid speed during forward movement. **C.** (Left) Representative kymographs of freely moving animals with AVA-specific constitutive optogenetic activation (+ATR). Controls (- ATR) were the same animals subjected to the same stimulation protocol but without the supply of the opsin activator. Right, Quantification of bending angles and centroid speed during forward movement. N: 14 (+ATR) and 10 (-ATR) animals. **D.** (Left) Representative kymograph of bending angles of animals subjected to AVA-specific optogenetic inhibition. (Right) Quantification of absolute bending angles and centroid speed during forward movement in non-stimulated (-ATR, N: 28) and stimulated (+ATR, N: 25) animals. **ATR**: all-trans retinal, co-factor for opsins expressed by transgenic *C. elegans*. Box-whisker plot displays the 90/10 and 75/25 percentiles, respectively. *: P < 0.05, **: P < 0.01, ***: P <0.001 by the Mann-Whitney U tests. P < 0.05 is considered to be statistically significant.

These findings establish an experimental paradigm of a bi-directional, fine-tuned manipulation of AVA’s excitability through its endogenous K+ leak.

### Depolarization of AVA potentiates both backward and forward movements

To determine the physiological consequence of AVA’s depolarized membrane potential, we applied CRISPR-Cas9 genome editing(*54*) and a repurposed ubiquitin-protease system-mediating protein degradation system(*55*) to achieve depletion of the endogenous TWK-40 activity specifically in the AVA neurons (Methods; Fig. 3S2). We address the role of AVA’s depolarized membrane potential, in context of an intact motor circuit and spontaneous locomotion.

In the absence of explicit sensory stimuli, a wild-type *C. elegans* spontaneously switches between forward and backward movements (*23*, *56*). Depleting TWK-40 from AVA alone produced overall increased motor activity (Movie 1). As expected, depleting TWK-40 from AVA increased the backward motor state: more frequent transitions into the backward motor state (Fig. 3S3A), with prolonged (Fig. 3S3A) and increased bending (Fig. 3S3A) during backward movement (Movie 1). Unexpectedly, depleting TWK-40 from AVA also potentiated forward locomotion: strikingly increasing bending during forward movement (Fig. 3B; Movie 1). These motor phenotypes were similar to *twk-40(lf)* genetic mutant animals (Fig 3S3B; Movie 1).

An increase in the animal’s overall motor activity upon an increased AVA membrane potential indicates AVA’s positive modulation of the forward circuit, instead of a strictly inhibitory role as proposed in the reciprocal inhibitory model.

Key support for the strictly reciprocal model was the previous observation that optogenetic depolarization of AVA resulted in instantaneous initiation and sustenance of backward movements (*57*, *58*). Indeed, in our strain that expresses an activating opsin (Chrimson) specifically in AVA (Methods), we recapitulated this result with strong and acute light stimulation (Fig. 3S4A; Movie 2).

A potential explanation for the discrepancy between the behavioral outcome of optogenetic and genetic depolarization of the AVA neuron is that the optogenetic stimulation protocol induced strong and acute depolarization that temporarily locks animals into the backward motor state. To address this possibility, we designed a different optogenetic stimulation protocol that more closely mimics genetic - fine-tuned and constitute - membrane depolarization (Methods). Indeed, constitute and moderate optogenetic activation of AVA potentiated forward movement (Fig. 3C; Movie 2).

### Hyperpolarization of AVA decreases both backward and forward movements

Mirroring our observation with the AVA-specific reduction of K+ leak, AVA-specific increase of K+ leak (AVA-TWK-40(gf)) (Methods) exerts a potent inhibitory effect on both forward (Fig. 3B) and backward (Fig. 3S3A) motor states, a behavior similar to that of *twk-40(gf)* mutant animals (Fig. 3S3A, B; Movie 1).

AVA-specific TWK-40(gf) and *twk-40(gf)* mutants not only exhibit drastically reduced overall motility, but also shared the characteristics where the backward motor state exhibiting slightly higher sensitivity: forward movement was drastically reduced (Fig. 3B; Fig. 3S3B) when backward movement was already fully abolished (Fig. 3S3A).

These defects could not be attributed, at least not fully to a potential developmental effect of constitutive hyperpolarization. When applying an inhibitory opsin (GtACR)- induced, AVA-specific acute silencing (Methods), we also observed strong inhibition of both the forward (Fig. 3D; Movie 3) and backward (Fig. 3S4B) movement.

Together, these results implicate that AVA’s depolarized membrane potential sustains the animal’s motility, during not only backward but also forward movements.

### AVA unidirectionally regulates AVB during spontaneous forward movement

AVA has no direct synaptic input to the major motor neuron classes that execute forward movement (*25*, *59*, *60*). To investigate the circuit mechanism that underlies AVA’s potentiation of forward movement, we examined the effect of ablating other interneurons that potentiate the executive motor neurons for forward movement (AVB, PVC) (*29*, *61*), or partition the forward and backward motor states (RIM) (*62*–*67*). PVC and RIM neurons are of further interest because they form stereotypic anatomic wiring with both AVA and AVB (Fig. 4S1A) (*24*, *68*), thus might facilitate indirect AVA-AVB communication.

We found that the ablation of AVB (Fig. 4S1B, Movie 4), but not of PVC or RIM (Fig. 4S1A; Movie 5), significantly attenuated the spontaneous forward movement of wildtype animals and *twk-40(lf)* mutants. Importantly, ablating AVB in AVA-specific TWK-40(lf) mutants also significantly reduced their forward movement (Fig. 4S1C; Movie 4).

These results concurred with conclusions from previous studies that AVB has a dominant role in potentiating the executive motor neurons for forward movement (*29*, *58*, *69*).

These results further indicate a potentially direct potentiation from AVA to AVB. This was unexpected because of the lack of stereotypic anatomic wiring from AVA to AVB (*24*, *68*).

To directly assessed this possibility, we examined the effect of AVA perturbation on AVB’s membrane potential through in situ electrophysiological recording (Fig. 4C, D). When AVA became further depolarized, AVB exhibited a significant increase in the amplitude of its potential changes (Fig. 4C; AVA-TWK-40(lf)) and a more depolarized membrane potential (∼-50mV to -10mV, median -32mV) (Fig. 4D). Conversely, when AVA became hyperpolarized, AVB’s potential change was drastically reduced in amplitude (Fig. 4C; AVA-TWK-40(gf)), with a modest hyperpolarization (∼-75mV to - 30mV, median -58mV) (Fig. 4D).

**Figure 4.**
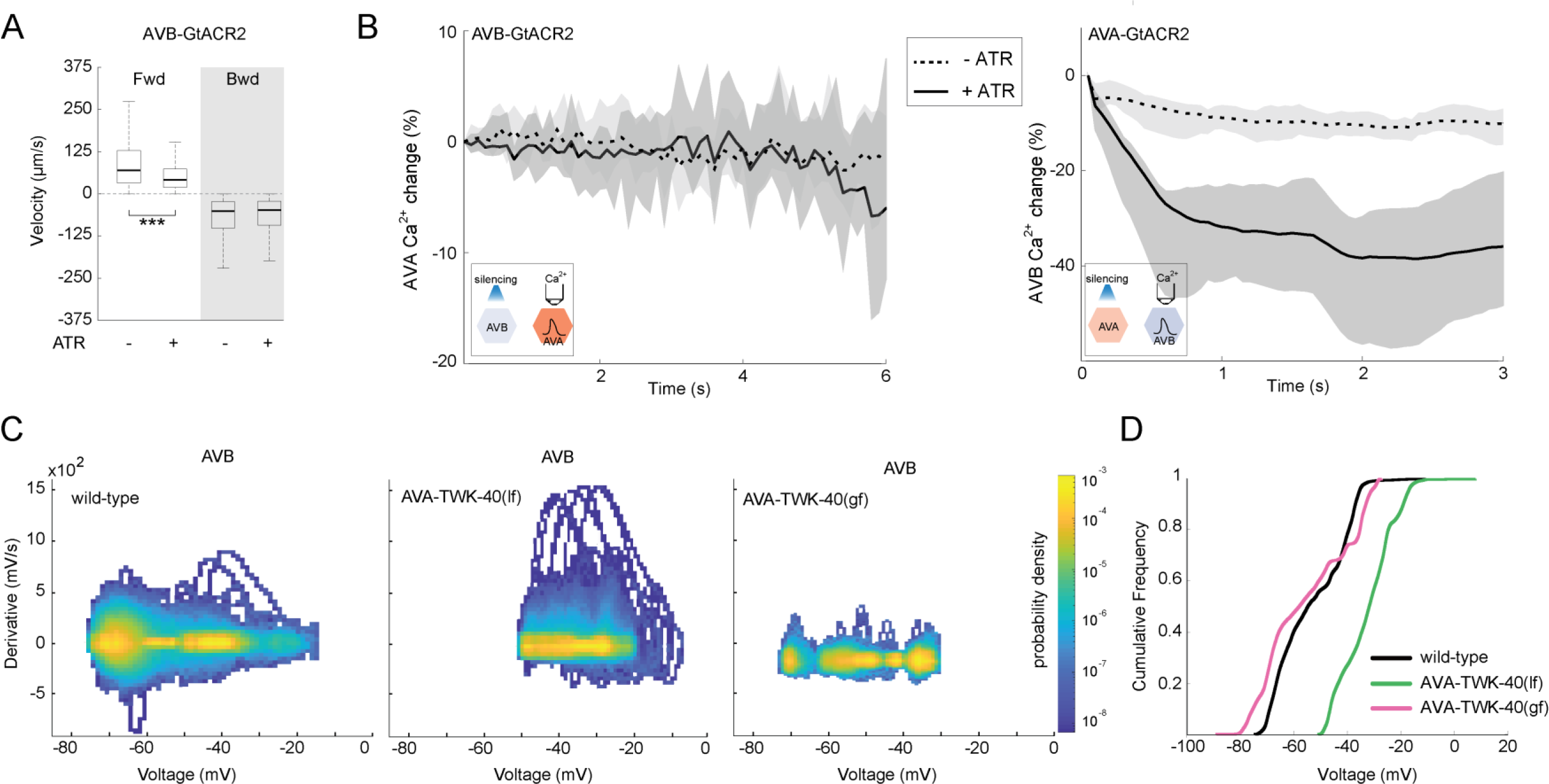
AVA potentiation of AVB is largely unidirectional. **A.** Quantification of mean velocity upon optogenetic inhibition of AVB by opsin (GtACR2). Silencing AVB led to decreased forward speed without affecting backward speed (ATR+; N: 11) compared to non-stimulated animals (-ATR; N: 5). **B.** (Left) AVA calcium activity in immobilized animals upon optogenetic inhibition of AVB by GtACR2. Inactivation of AVB (+ATR; N: 28) did not lead to consistent changes in AVA calcium dynamics compared to un-stimulated animals (-ATR; N: 19). (Right) AVB calcium activity in immobilized animals upon optogenetic inhibition of AVA. Inactivation of AVA (+ATR; N: 23) led to reduced AVB calcium dynamics compared to un-stimulated animals (-ATR; N: 23). Lines and shaded bars denote the median and 95% confidence interval, respectively. Insert panels illustrate the experimental setup. **ATR**: all-trans retinal, co-factor for opsins expressed by transgenic *C. elegans*. **C.** Distribution of the voltage derivative across the AVB membrane potential range and dynamics in respective genotypes. In AVA-specific TWK-40(*lf)* (N: 7 cells) and AVA-specific TWK-40(*gf*) (N: 7 cells) mutant animals, AVB exhibited an increased and decreased voltage dynamic range, respectively, compared to wild-type animals (N: 8 cells). **D.** Cumulative curves of the AVB membrane potential range and dynamics, where the AVB neuron exhibited a more depolarized and hyperpolarized potential range in AVA-specific *twk-40(lf)* (N: 7 cells) and AVA-specific *twk-40(gf)* (N: 7 cells) mutant animals, respectively, compared to wild-type animals (N: 7 cells). Each AVB neuron was recorded for 60-200 seconds.

We asked whether this effect is reciprocal. Specifically, whether AVB could also potentiate AVA. We examined the effect of AVB inhibition on the backward movement. As expected, optogenetic silencing of AVB reduced the forward movement in freely moving animals (Fig. 4A), and decreased AVB’s calcium activity in immobilized animals (Fig. 4S2A). In these animals, the backward movement was unchanged (Fig. 4A), and AVA’s activity pattern was similar with or without AVB silencing (Fig. 4B).

The lack of a significant effect of AVB silencing on AVA contrasted the effect of AVA silencing on AVB (Fig. 4D). Consistent with our finding that silencing AVA activity led to attenuated forward movement (Fig. 3), AVA-specific optogenetic silencing reduced the calcium activity in not only AVA (Fig. 4S2B), but also AVB (Fig. 4B) in immobilized animals.

These experimental findings reveal an asymmetry between the forward motor circuit and backward motor circuit. Reflected by the relationship between AVA and AVB, communication is predominantly from AVA to AVB, instead of a strict reciprocal inhibition, during spontaneous locomotion.

### A two-timescale model predicts AVA’s regulation of AVB

These results raised the need to directly examine the relationship between AVA and AVB neuronal activity, which has not been simultaneously analyzed in freely moving animals in previous studies (*5*, *70*, *71*).

We performed simultaneous calcium imaging of AVA and AVB during spontaneous locomotion (Methods). For both neurons, the largest activity transients occurred during motor state transitions. As reported (*28*, *29*, *70*, *72*–*76*), each epoch of AVA transients aligned with a transition event. Specifically, the initiation of a backward movement coincided with the onset of AVA’s calcium rise, whereas the exit from a backward movement coincided with the onset of AVA’s calcium decay (Fig. 5SA, left panels).

During this period, AVB’s calcium change was anti-correlated with that of AVA (Fig. 5SB, left panel).

However, this relationship changed after animals began an exit from the backward movement. While AVA’s activity rapidly decayed to a steady baseline, AVB showed a continued rise as the animal exited the backward movement, followed by a slow decay after the animal entered forward movement (Fig. 5SA, left panels). During subsequent forward movement, AVA maintained its baseline calcium signals (Fig. 5SA, left panels).

We sought the simplest mathematical model to describe this dynamic relationship. Because the functional communication between these two neurons was largely unidirectional during spontaneous locomotion (Fig. 4; Fig. 4S2), we considered an input-output model where AVA is treated as the input to AVB. The calcium imaging data (Fig. 5A; Fig. 5SA) suggested a model with two timescales: a fast input during backward movements, and a slower one during post-backward movements, to account for two phases of AVB activity dynamics. This led to a simple linear dynamical system model (Fig. 5B).

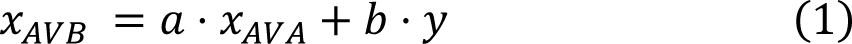

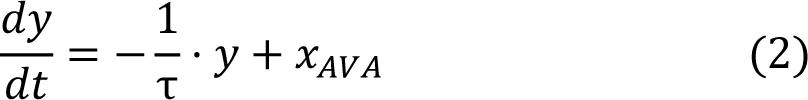

where *x_AVB_* and *x_AVA_* represent the activity of AVB and AVA, respectively; *y* represents the slow input from AVA (1), given by the leaky-integrator in (2).

**Figure 5.**
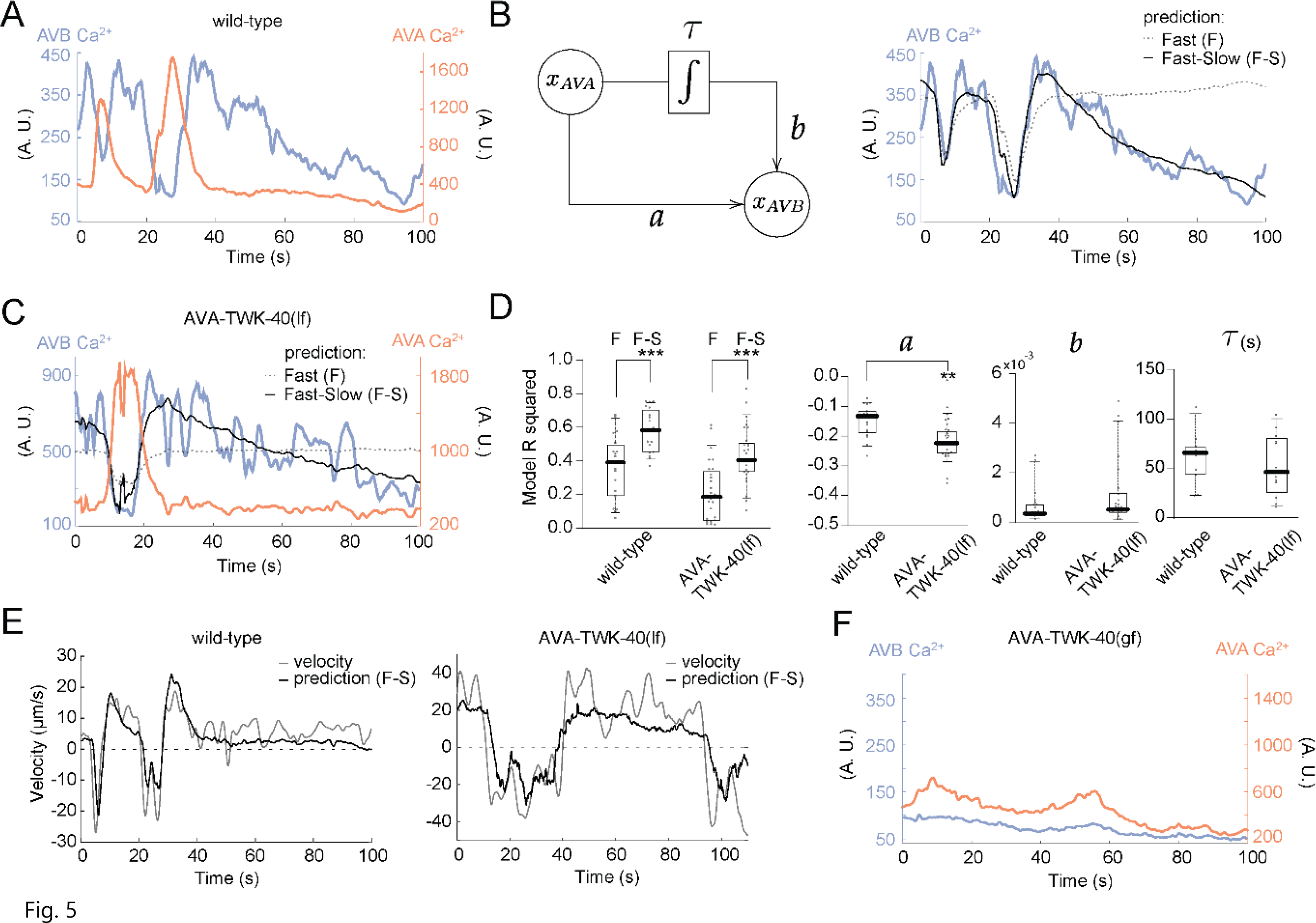
A two-timescale model predicts AVA and AVB’s relationship. **A.** Representative calcium imaging traces of co-imaged AVA and AVB during spontaneous movements by a wild-type animal. **B.** (Left) AVB activity is modeled as a sum of two terms dependent on AVA’s activity. A fast copy with a proportionality constant *a* and a slow copy integrated with a timescale τ and proportionality constant *b*. (Right) The fast-slow model is a better fit for AVB’s post-backward movement activity compared to a model without the slow timescale. **C.** Example calcium traces of AVA and AVB from AVA-specific TWK-40(lf) mutant animals show that the fast-slow model also explains AVB’s activity when AVA was further depolarized. **D.** (Left panel) Model R- squared values comparing the fast-slow model with the fast only models, for both wild-type and AVA-specific-TWK-40(lf) animals. Removing the slow time scale input decreases model performance in both cases. (Right panels) Comparison of the estimated value for parameters for *a*, *b,* and τ (the integration timescale for the slow copy) between wild-type and AVA-specific TWK-40(lf) traces. Biologically, the two-timescale model suggests a fast inhibition (negative *a*) and a slow excitation (positive *b*) from AVA to AVB. N: wild-type - 11 epochs of backward movement from 11 animals; AVA-specific TWK-40(lf) - 10 epochs of backward movement from 10 animals. Box-whisker plot displayed the 90/10 and 75/25 percentiles, respectively. *: P < 0.05, **: P <0.01, ***: P <0.001 by the Mann-Whitney U tests. P < 0.05 is considered to be statistically significant. **E.** Sample traces of the F-S model prediction of the velocity of (left) wild-type animals and (right) AVA-specific TWK-40(lf) mutant animals from AVA calcium activity. Parameters: wild-type *a* = −0.51,*b* = 0.02, τ = 3.5s; AVA-specific TWK-40(gf): *a* = −0.38, *b* = 0.004, τ = 16.6s. **F.** Representative calcium activity traces of co-imaged AVA and AVB by a freely moving AVA-Specific-TWK-40(gf) mutant animal. Both neurons showed strong reduction in activity.

This model has three free parameters: the strength of the fast input, *τ*, the strength of the slow input, *b*, and the timescale for the slow input, τ. We used the framework of linear systems identification to fit these parameters to our experimental calcium imaging data (Fig. 5C; Methods). The best-fit parameters set the strength of fast input *a* to a negative value, and the slow input *b* to a positive value (Fig. 5D, wild-type).

Biologically, this could be interpreted as the forward motor circuit receiving inputs from AVA at two timescales with opposite polarities: a fast inhibitory input and a slow excitatory input. In other words, AVA maintains a constitutive and excitatory input on AVB, in addition to a strong, but activation-mediated phasic inhibitory input to AVB during transitions. Without the slow excitatory input, the one timescale model cannot predict the gradual decay of AVB activity after a backward movement (Fig. 5B, D, wild-type).

With different parameter values, this two-timescale model could also predict the animal’s spontaneous velocity from AVA’s activity (Fig. 5E, wild-type), thus capturing the continuous modulation of behavior during spontaneous state transitions.

Together, our experimental data and data-driven modelling leads to a revised model, where AVA functions as a master regulator of spontaneous locomotion by exerting bi-directional regulation of AVB activity at different timescales. A transient but strong depolarization of AVA leads to phasic inhibition during backward movements, while a persistent but weak depolarization of AVA provides tonic excitation to AVB that promotes forward movement between backward movements.

### Depolarized membrane potential is necessary for AVA to function as a master regulator

This model led us to ask if AVA’s depolarized membrane potential is required for this master regulator role. We co-imaged AVA and AVB calcium activity in freely behaving AVA-specific TWK-40(lf) and AVA-specific TWK-40(gf) mutants, and examined the effect of altered AVA membrane potential on their relationships.

We found that with a further membrane depolarization (AVA-TWK-40(lf)), AVA’s time-dependent relationships with AVB neuron and animal’s spontaneous velocity are maintained (Fig. 5SA, middle panels; Fig. 5B, AVA-TWK-40(lf)). The two-timescale model was sufficient to predict AVB calcium activity (Fig. 5C, D) and velocity (Fig. 5E, AVA-TWK-40(lf)) from AVA’s calcium signals. Parameters for both inputs showed a trend of further increase, but with a slightly higher value for the fast input *a* (Fig. 5D, AVA-TWK-40(lf)).

Biologically, this suggests that with further depolarization, AVA maintains and might strengthen both fast-inhibitory and slow-excitatory inputs. This is consistent with the behavior – a potentiation of both forward and backward movement – of these animals.

In AVA-TWK-40(gf) mutants, the relationship between AVA and AVB activity changed completely such that a model was no longer applicable. Reflecting the attenuation of locomotion upon hyperpolarization of AVA membrane potential, both neurons exhibited drastically reduced calcium signals with little dynamical changes (Fig. 5F; Fig. 5SA, right panels).

These results suggest that the two-timescale model for AVA’s master regulator role is contingent on its depolarized membrane potential.

### Cholinergic transmission plays a major role in AVA’s tonic excitation of the forward motor state

AVA’s constitutive input onto the forward circuit, which depends on a maintained depolarized membrane potential, implicates its tonic activation.

We further evaluated this possibility with a dosage-dependent chemogenetic hyperpolarization by a histamine-gated chloride channel (HisCl) (*36*). We reproduced results from the previous study: exposing animals that ectopically express HisCl in AVA led to a histamine-dependent reduction of backward speed, as well as spontaneous forward speed (*36*). We further noted that the backward motor circuit exhibited higher sensitivity to histamine. While the backward movement was fully blocked at the presence of 2mM histamine (Fig. 6A, left panel), the forward movement was only modestly reduced (Fig. 6A, middle panel). With increased histamine concentration, the forward movement progressively declined (Fig. 6A, middle panel).

**Figure 6.**
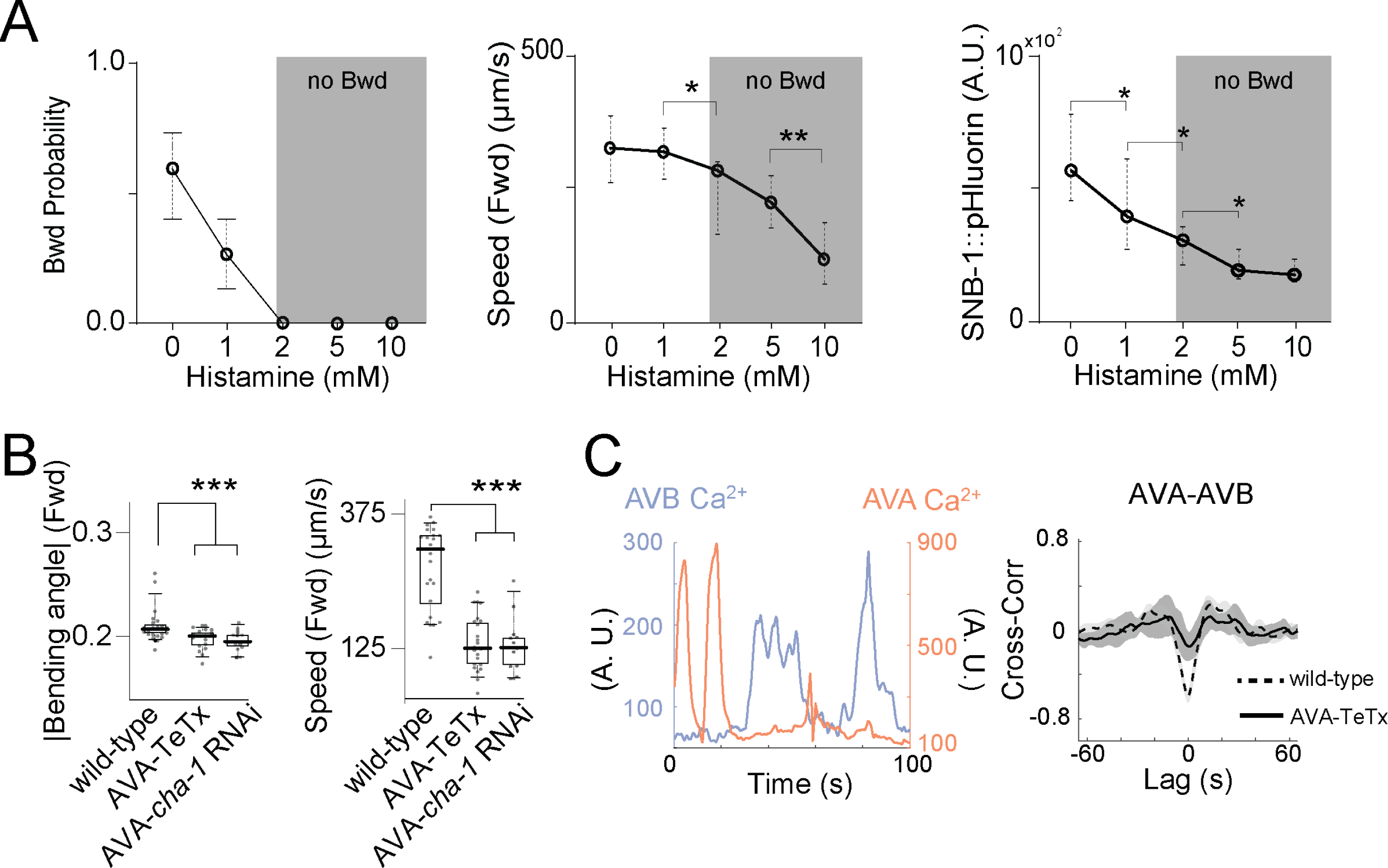
Tonic acetylcholine release from AVA potentiates forward movement. **A.** (Left) Backward movement probability upon inhibition of AVA by HisCl at different histamine concentrations. Histamine impaired spontaneous backward movement in a dosage-dependent manner. N: 15 animals at each concentration. (Middle) Centroid forward speed upon inhibition of AVA by HisCl at different histamine concentrations. The forward speed exhibited a gradual reduction upon increased histamine. N: 15 animals per concentration. (Right) Mean AVA-specific SNB-1::pHluorin signals upon inhibition of AVA by HisCl at different histamine concentrations. While showed a continuous decrease with increased histamine concentration. While the spontaneous backward movement was already fully blocked at 2mM histamine, fluorescent signals persisted, and continuously decrease with increased histamine. N: 30 (0mM), 29 (1mM), 30 (2mM), 31 (5mM), 28 (10 mM) animals. Shaded areas denote Histamine’s concentration range where we observed complete abolishment of spontaneous backward movement. Quantification showed median and 95% confidence interval. **B.** Mean absolute bending angles (Left panel) and mean centroid speed (Right panel) during forward movement by animals with AVA’s synaptic vesicle fusion blocked (AVA- TeTx), or vesicular acetylcholine loading blocked (AVA-*cha-1* RNAi). Forward movements were reduced by AVA-specific TeTx and AVA-*cha-1* RNAi. N: 23 animals (Control), 18 animals (AVA-specific TeTx), 14 animals (AVA-*cha-1* RNAi). Box-whisker plot displays the 90/10 and 75/25 percentiles, respectively. **C.** (Left panel) A representative traces of co-imaged AVA and AVB calcium activity in a freely moving AVA-specific TeTx animal. (Right panel) Mean cross-correlations of co-imaged AVA and AVB activity in wild-type and AVA-specific TeTx animals. Positive and negative correlation with the backward speed and post-backward forward speed, respectively, exhibited by wild-type animals, were both reduced in AVA-specific TeTx animals. N: 32 epochs from 10 AVA-specific TeTx animals; 21 epochs from 11 wild-type animals. *: P < 0.05, **: P < 0.01, ***: P <0.001 by the Mann-Whitney U tests. P <0.05 is considered to be statistically significant.

An implication of such behavioral response is that with sequential membrane hyperpolarization, AVA might rapidly lose its ability of transient activation (which underlies its phasic inhibitory input onto the forward circuit), but gradually lose its constitutive activation (which underlies its excitatory input onto the forward circuit), thus separating tonic excitation and phasic inhibition of the forward movement.

Consistent with AVA being tonically excitatory, synaptic vesicle exocytosis from AVA, visualized by SNB-1::pHluorin, was reduced but remained at 2mM histamine when the phasic inhibition was fully blocked (Fig. 6A, right panel B). In alignment with SNB-1::pHluorin signal level reflecting AVA activity, depolarization and hyperpolarization of AVA by *twk-40(lf)* and *twk-40(gf)* genetic mutations led to a respective increase and decrease of SNB-1::pHluorin signal from the AVA neuron (Fig. S61A).

Lastly, we asked whether AVA’s constitutive inputs to the forward motor circuit requires acetylcholine, as AVA functions primarily as a cholinergic neuron but also synthesizes many neuropeptides (*77*–*79*). Tetanus toxin (TeTx) potently blocked synaptic vesicle-mediated transmission in AVA (Fig. 6S2A), but much less so with dense core vesicle-mediated transmission (Fig. 6S2B-D). AVA-specific expression of TeTx (AVA-TeTx) led to a significant decrease in spontaneous forward movements, in both wild-type animals (Fig. 6B, AVA-TeTx) and *twk-40(lf)* mutants (Fig. 6S1B, AVA-TeTx). AVA-TeTx fully also fully blocked AVA-mediated, optogenetic potentiation of forward movement (Fig. 6S1C).

These results indicate that AVA’s potentiation of the forward motor circuit might primarily depend on synaptic vesicle-mediated transmission. This was corroborated by the effect of reducing vesicular loading of acetylcholine, through AVA-specific RNA interference(*80*) against the acetyltransferase CHA-1 (*81*) (Methods). AVA-*cha-1* RNAi led to a characteristically similar, but more moderate reduction of forward movement when compared to AVA-TeTx (Fig. 6B).

Consistent with its behavioral effect, AVA-TeTx disrupted AVA’s relationships with AVB activity (Fig. 6C). In AVA-TeTx animals, AVA continued to exhibit calcium transients (Fig. 6C). However, the correlation between AVA and AVB calcium dynamics, during both the positive and negative phases, was attenuated (Fig. 6C, D).

We conclude that the tonically active premotor interneuron AVA functions as a master neuron of the *C. elegans* motor circuit. AVA contributes to both constitutive activation and transient inhibition of the forward motor circuit, which requires cholinergic signaling. Together with our previous findings on AVA’s dual regulation of the backward motor circuit (*27*, *28*, *82*), these results lead to an elegant mechanistic model that underlies a spontaneous, continuous motility consisted of dynamic transitions between two mutually opposing motor states (Fig. 7), to be elaborated in Discussion.

**Figure 7.**
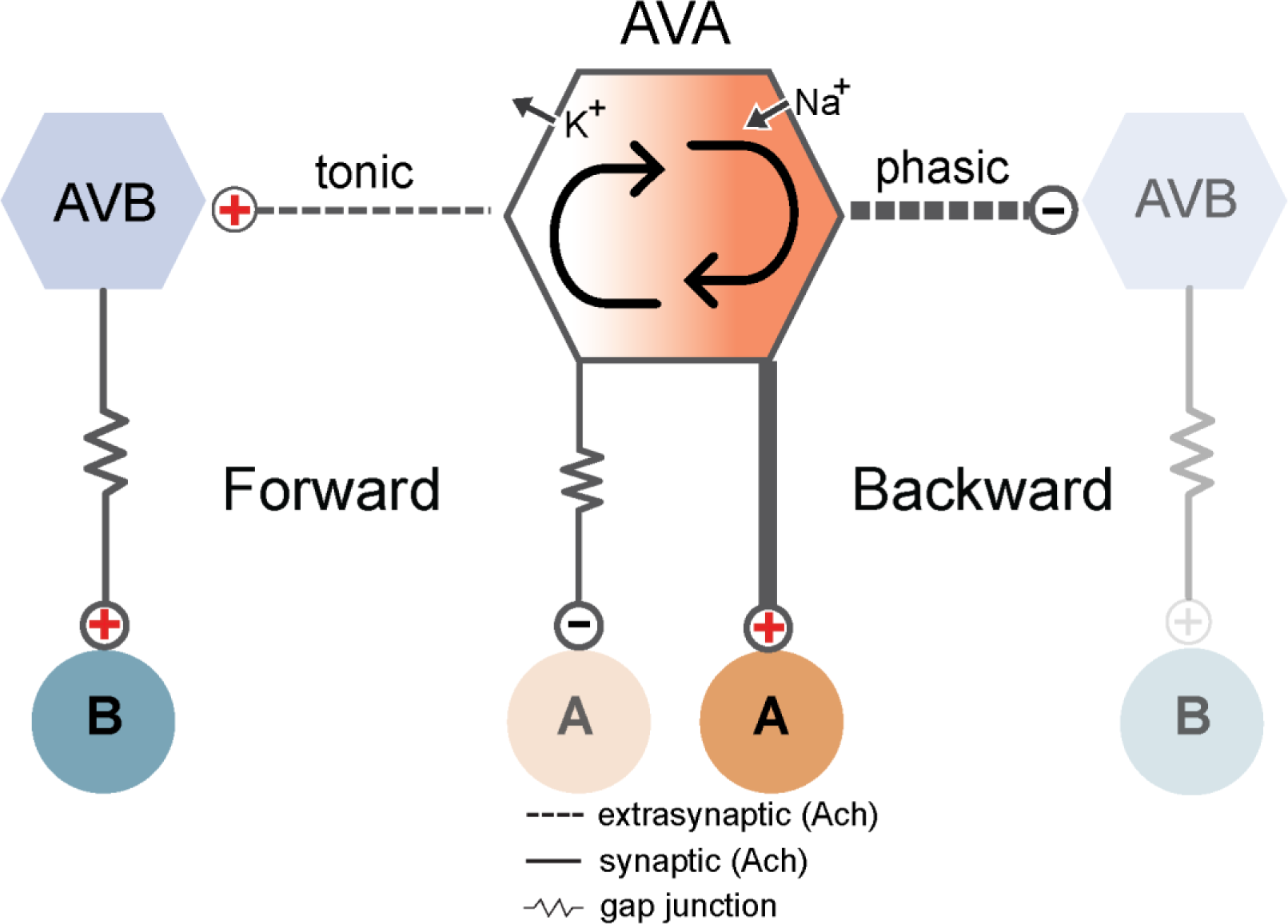
A master neuron configuration for continuous modulation and transitions between directional movements. AVA functions as a master regulator to coordinate the forward and backward movement and their smooth transitions. This modulatory neuron depends on cyclic changes of AVA activity. **(Left)** At its lower activity state, AVA actively shunts the backward motor neurons A by gap junction-coupling(*71*). At this state, AVA’s tonic potentiation of AVB, which activates the forward motor neurons B(*29*, *69*), dominates. This ensures sustained forward movement. **(Right)** At its high activity state, AVA phasically activates the backward motor neurons A by chemical synaptic transmission(*71*). At this state, AVA exerts phasic inhibition of AVB, which initiates and sustains the backward movement. Arrows between the low and high AVA states denote continuous modulations of AVA activity during transitions between the mutually exclusive forward and backward motor states. Dashed and solid lines denote extrasynaptic (indirect) and synaptic connectivity, respectively; thickness denotes communication strength.

## Discussion

Locomotion is composed of continuous modulation of different behavioral motifs and transitions between them (*1*–*11*). Here we address how a neural system enables this continuous modulation and transition. We summarize below that in *C. elegans*, a tonically active premotor interneuron serves this role, through a bi-directional regulation of two opposing motor states in a master-slave configuration. This revised model offers several potentially general biological and evolutionary implications, described after introducing the model.

### A master-slave model for continuous motor behaviors

*C. elegans* intrinsically switches between the forward and backward motor states(*23*). Previously, we found that AVA exerts dual modulation of the backward circuit(*27*, *28*, *82*): at ‘rest’, AVA reduces spontaneous backward movement through gap junctional-mediated shunting motor neurons that execute backward movement, but initiates backward movement by activating the same motor neurons with chemical synapses when ‘activated’ (Fig. 7, center panel). Our new finding - AVA’s chemical synapse also modulates the forward circuit, with opposite polarities at different time scales (Fig. 7, side panels) - reveals the other side of the same coin. Together, it offers a parsimonious circuit solution for overall motor control by a single neuron.

In this model, AVA is tonically active but switches between a high and low activity state. At its high state, its chemical synapse simultaneously excites the backward circuit and inhibits the forward circuit, which leads to backward movement. As AVA’s activity decreases, its chemical synapse’s sustained excitation of the forward circuit and its gap junction’s sustained inhibition of the backward circuit prevail, leading to the exit of backward movement and transition into the forward movement.

This model makes AVA the master neuron that continuously regulates two opposing motor motifs, mediating their intrinsic transitions without risking losing overall motility.

### Asymmetry between the forward and backward motor states

This configuration breaks an assumed functional symmetry between the forward and backward motor circuits.

An inherent asymmetry between the forward and backward functional modules explains several long-noted behavioral characteristics. During spontaneous locomotion, *C. elegans* preferentially sustains forward movement, resulting in a higher propensity of the forward state (*23*). But when executed, the shorter backward movement exhibits greater vigor than the forward movement(*83*). Backward movement is more frequently evoked and prolonged when animals encounter repulsive stimuli or attempt to escape (*23*, *83*, *84*). Importantly, evoked backward movement potently modifies the forward state that follows. A prolonged backward movement is more likely followed by a turn, a form of forward movement that steers away from the original trajectory, whereas a shorter backward movement is more likely followed by forward movement in a similar trajectory (*23*, *56*, *85*, *86*), which resembles most spontaneous backward movement. Evoked backward movement further increases the forward speed, potentiating the forward state for up to several minutes (*34*, *35*). Backward movement thus represents an intrinsically more robust, but actively suppressed motor state. Its activation modifies the subsequent forward state.

### Anatomic wiring does not account for AVA’s full functional connectivity

AVA’s function during spontaneous locomotion only partially correlates with its anatomical wiring (*24*, *59*, *68*).

Anatomically, AVA makes extensive chemical and electrical wiring with motor neurons that execute backward movement (*24*, *59*, *68*). This synaptic connectivity correlates with AVA’s regulation of the backward circuit (*27*, *28*, *32*, *82*, *87*–*89*).

In contrast, AVA’s modulation of the forward circuit cannot be predicted by its anatomical wiring. AVA tonically activates AVB, which contributes to not only sustained forward movement, but also inherently brief backward movement. However, AVA does not have direct synaptic inputs to AVB, whereas AVB makes direct inputs to AVA (Fig. 1A) (*24*, *68*).

AVA’s tonic excitation requires its depolarized membrane potential and its cholinergic synaptic transmission. Our results suggest a mechanism in which its depolarized membrane allows constitutive acetylcholine release as a facilitation of extra-synaptic accumulation and potentiation of the forward motor circuit. Indeed, in the developing larvae, extra-synaptic cholinergic transmission from premotor interneurons provides the tonic potentiation of ventral muscles that functionally substitute for the entire missing ventral subcircuit (*90*).

This study joins an expanding report on the lack of a straight-forward correlation between anatomic wiring, population neuronal activity, and behaviors, across systems (*91*–*95*). Significant inconsistency between functional and anatomical neuronal connectivity implies the necessity of studying other forms of functional communication - ephaptic coupling, extrasynaptic transmission, and modulatory signaling (*91*–*94*).

A further, perhaps more interesting intuition from our study is that in a small neural system, with limited numbers of anatomic wiring, the form of ‘invisible’ wiring might be preferentially utilized for spontaneous behavioral states, whereas anatomic wirings, such as AVB to AVA, is built mainly for contextual recruitment for decision-making or evoked behaviors.Essentially, the functional-anatomic connectivity discrepancy might reflect the extend of economical wiring, as the size and cellular complexity of neural systems evolve.

### A parsimonious role of neurons of the backward motor circuit

Behaviorally*, C. elegans* spends most of its time in the forward state. It is thus intriguing that AVA, the regulator of overall motor control, is the premotor interneuron that is anatomically wired to gate motor neurons that execute the backward movement.

We posit that a motor circuit might be built first to generate robust escape responses, which are crucial for survival. The escape circuit is then actively suppressed and repurposed for forward locomotion. In other words, the motor system finds an economical solution by using the same tonic drive to shunt backward and potentiate forward locomotion. Its tonic excitatory input to the forward circuit, an integrated history of its activity, might further serve as the short-term memory of the internal states and environment.

### Dynamic regulation of a non-spiking neural network b membrane potential

With a few exceptions (*95*, *96*), the *C. elegans* neural system consists primarily of non-spiking neurons (*39*, *43*, *44*, *97*–*101*). This neural network generates behaviors that are stable, flexible, and adaptive, as those that function through regenerative potentials.

AVA’s functional model suggests that precise and dynamic regulation of the activity of non-spiking neurons and a network consisted primarily by non-spiking neurons can be achieved through the precise modulation of individual membrane potentials.

When a neuron functions through graded potential changes, it is particularly suited for a K2P conductance to regulate its activity range and form of responses to stimuli(*102*, *103*). *C. elegans* exceptional expansion of K+ channels, in particular, the 47 K+ leak channels (*52*, *103*–*105*) is consistent with this role. With 47 K2P genes expressed by overlapping subgroups of 300 neurons (*78*), these channels could define the unique computational properties of each neuron.

### Unsolved questions and future studies

We have focused on spontaneous locomotion. In particular, our model is intended to represent the simplest possible dynamical system that explains AVA-AVB calcium recordings during spontaneous locomotion. Reassuringly, all the experimental data in this study is consistent with our model. However, this study does not exclude more complex mechanisms, for example, extensive reciprocal feedback between AVA and AVB, non-cholinergic transmission, and indirect communications via other neurons (such as RIM and PVC), to be involved during goal-oriented navigation and escape responses. In fact, external stimuli and adaptive behaviors change not only the ratio, but also the form of motor state transition by a probabilistic addition of a turning module (*56*, *106*), a feature that we did not attempt to model.

We revealed the importance of cholinergic transmission from AVA, without knowing the identity of genes that underlie its effects on AVB. This is not a trivial task. The *C. elegans* nervous system possesses a large number of cholinergic receptors that belong to multiple classes and exhibit significant functional redundancy (*107*). Similar cholinergic receptor complexes are re-iteratively applied, present in interneurons, excitatory motor neurons, inhibitory motor neurons, as well as body wall muscles (*78*, *107*). Together, they raise considerable technical challenges towards pinpointing a cell-specific role. Multiple dedicated future studies would be required to address this issue.

**Table 1:** Membrane potentials (MP) of *C. elegans* neurons.

| Neuron class | Neuron | Approximate 'Rest' MP |
| --- | --- | --- |
| Sensory neuron | AFD | -40 mV(101, 110), -80 mV(95) |
| Sensory neuron | AWA | -75 mV(95, 111) |
| Sensory neuron | PLM | -60 mV(100) |
| Sensory neuron | ASE | -55 mV(39) -60mV(112) |
| Sensory neuron | ASH | -60 mV(98) |
| Interneuron | AIY | -55 mV(95) |
| Interneuron | AIA | -80 mV(113) |
| Premotor interneuron | AVA | -20 to -30 mV(38, 42, 43) |
| Premotor interneuron | AVB | -75 to -55 mV (this study) |
| Premotor interneuron | AVE | -35 mV (S. Gao, unpublished) |
| *A sra-11* expressed neuron | unsure | -20 mV(114) |
| Motor neuron | RMD | -70 mV and -35 mV(43) |
| Motor neuron | RIM | -40 mV(95) |
| Enteric motor neuron | DVB | -60 mV(96) |
| Enteric motor neuron | AVL | -40 mV(96) |
| Motor neuron | VA5 | -70 mV(44) |
| Motor neuron | VB6 | -50 mV(44) |
| Motor neuron | VD5 | -45 mV(44) |

**Table 2.** Strains and constructs generated and/or used in this study.

| Strain | Genotype | Plasmid | Description (see Notes) | Figures |
| --- | --- | --- | --- | --- |
| **Electrophysiology recording (visualization of AVA and AVB)** |  |  |  |  |
| ZM11177 | *hpls860* | pJH4693 | [*Ptwk-40s-eGFP; \*Pmyo-2-wCherry*] | 1B-D; 3A; 3S1 |
| ZM11207 | *twk-40(bln336); hpls860* | pJH4693 | *twk-40(gf); [Ptwk-40s-eGFP; \*Pmyo-2-wCherry]* | 3A; 3S1 |
| ZM11208 | *twk-40(hp834); hpls860* | pJH4693 | *twk-40(lf); [Ptwk-40s-eGFP; \*Pmyo-2-wCherry]* | 3A; 3S1 |
| ZM11145 | *hpls768/+; hpEx4405* | pJH4316<br>pJH4237<br>pJH4693 | <u>AVA-specific TWK-40(gf)</u><br>[*Pf1p-18LoxPstopLoxPtwk-40(L202N)::GFP Pnmr-1-FLP*];<br>[*Ptwk-40s-eGFP; \*Pmyo-2-wCherry*] | 4C, D |
| ZM11139 | *twk-40(bln282) hpEx3904; hpEx4401* | pJH4018<br>pJH2673<br>pJH4693 | <u>AVA-specific TWK-40(lf)</u><br>*twk-40::TagRFP::ZF*<br>[*Prig-3-FRT-stop FRT twk-40(gf); Ptwk-40s-FLP, \*HygR*]<br>[*Ptwk-40s-eGFP; \*Pmyo-2-wCherry*] | 4C, D |
| ***twk-40* mutants and locomotion characterization** |  |  |  |  |
| ZM8302 | *twk-40(hp834)* | CRISPR/<br>Cas9 | *twk-40(lf)* | 2B; 3S3 |
| JIP1514 | *twk-40(bln336)* | CRISPR/<br>Cas9 | *twk-40(gf)* | 2B; 3S3 |
| JIP1368 | *twk-40(bln282)* | CRISPR/<br>Cas9 | *twk-40::TagRFP::ZF* | 3S2 |
| ZM10113 | *twk-40(bln282); hpls727* | pJH3830 | <u>AVA-specific TWK-40(lf)</u><br>*twk-40::TagRFP::ZF*;<br>[*Prig-3-zif-1 sl2 RFP; \*Pttx-3-GFP*] | 3B; 3S3 |
| ZM10066 | *hpEx4053* | pJH4203<br>pJH2673 | <u>AVA-specific TWK-40(gf)</u><br>[*Prig-3-FRTstopFRT twk-40(L202N)::GFP Pnmr-1-FLP; \*Pmyo-2-wCherry*] | 3B; 3S3 |
| **Immunostaining** |  |  |  |  |
| ZM10355 | *twk-40(bln282); hpEx4120* | pJH1371 | *twk-40::TagRFP::ZF*;<br>[*Prgef-1-GPI::YFP*] | 2C |
| **PVC ablation** |  |  |  |  |
| ZM7054 | *hpls321* | pJH2827 | <u>AVA, AVE, AVD, RIM, PVC mini SOG</u><br>[*Pnmr-1-tomm-20::miniSOG sl2 wCherry; \*Lin-15(+)*] | 4S1A |
| **RIM ablation (RIM miniSOG)** |  |  |  |  |
| ZM7978 | *hpls327* | pJH2829 | [*Pcex-1-tomm-20::miniSOG sl2 wCherry*;<br>* *Lin-15(+)*] | 4S1A |
| **AVA-specific TeTx and *cha-1* RNAi** |  |  |  |  |
| ZM10755 | *hpls814* | pJH4508<br>pJH4237 | AVA-specific TeTx<br>[*Pf1p-18-LoxP EBFP stop LoxP TeTx::RFP* ;<br>*Ptwk-40s-Cre*] | 6B |
| ZM11082 | *twk-40 (hp834)* ;<br>*hpls814* | pJH4508<br>pJH4237 | *twk-40(lf)*;<br>[*Pf1p-18-LoxP EBFP stop LoxP TeTx::RFP*;<br>*Ptwk-40s-Cre*] | 6S2B |
| ZM11086 | *hpEx4362* | PCR<br>product | AVA-specific *cha-1* RNAi<br>[*Pf1p-18-cha-1(sense)*; *Ptwk-40-cha-1*<br>(antisense); * *Pttx-3-GFP*] | 6B |
| **AVA peptide secretion and synaptic vesicle analyses** |  |  |  |  |
| ZM11039 | *hpEx4351* | pJH4677 | [*Pnpr-4-ins-22::GFP*] | 6S1C, D |
| ZM11078 | *hpEx4351*;<br>*hpEx4364* | pJH4677<br>pJH4506 | with AVA-TeTx<br>[*Pnpr-4-ins-22::GFP*];<br>[*Pnpr-4-TeTx-wCherry*] | 6S1C, D |
| ZM11084 | *hpEx4363* | pJH4687 | [*Pnpr-4-ins-22::pHluorin*; * *Pmyo-2-wCherry*] | 6S1B |
| ZM11085 | *hpEx4363*;<br>*hpls814* | pJH4687<br>pJH4508<br>pJH4237 | with AVA-TeTx<br>[*Pnpr-4-ins-22::pHluorin*; * *Pmyo-2-wcherry*];<br>[*Pf1p-18-LoxP EBFP Stop LoxP TeTx::RFP*;<br>*Ptwk-40s-Cre*] | 6S1B |
| ZM10040 | *hpls721* | pJH4169<br>pJH2673 | AVA-specific vesicle marker<br>[*Prig-3 FRT stop FRT SNB-1::GFP*;<br>*Pnmr-1-FLP*; * *Pmyo-2-wcherry*] | 6S1A |
| ZM10786 | *hpls721*;<br>*hpls811* | pJH4169<br>pJH2673<br>pJH4508<br>pJH4237 | with AVA-TeTx<br>[*Prig-3 FRT stop FRT SNB-1::GFP*;<br>*Pnmr-1-FLP*; * *Pmyo-2-wCherry*];<br>[*Pf1p-18-LoxP EBFP Stop LoxP TeTx::RFP*;<br>*Ptwk-40s-Cre*] | 6S1A |
| ZM11287 | *hpEx4479* | pJH4783 | [*Pnpr-4-SNB-1::pHluorin*; * *Lin-15*] | 6S2A |
| ZM11284 | *twk-40(bln336)*;<br>*hpEx4479* | pJH4783 | *twk-40(gf)*;<br>[*Pnpr-4-SNB-1::pHluorin*; * *Lin-15*] | 6S2A |
| ZM11285 | *twk-40 (hp834)*;<br>*hpEx4479* | pJH4783 | *twk-40(lf)*;<br>[*Pnpr-4-SNB-1::pHluorin*; * *Lin-15*] | 6S2A |
| **AVA optogenetic (Chrimson) activation** |  |  |  |  |
| ZM10206 | *hpls758* | pJH4253<br>pJH4237 | [*Prig-3 LoxP EBFP stop LoxP Chrimson::RFP*;<br>*Ptwk-40s-Cre*] | 3B; 3S4A |
| ZM11151 | *hpls758*;<br>*hpls814* | pJH4253<br>pJH4237<br>pJH4508<br>pJH4237 | with AVA-TeTx<br>[*Prig-3-LoxP EBFP stop LoxP Chrimson::RFP*;<br>*Ptwk-40s-Cre*];<br>[*Pf1p-18-LoxP EBFP stop LoxP TeTx::RFP*;<br>*Ptwk-40s-Cre*] | 6S2C |

| **AVA HisCl chemogenetic inactivation** |  |  |  |  |
| --- | --- | --- | --- | --- |
| ZM9354 | *hpls636* | Ref.(36) | *[Prig-3::HisCl1::SL2::mCherry]* | 6A, left/mid |
| ZM11286 | *hpls636*;<br>*hpEx4479* | Ref.(36)<br>pJH4783 | *[Prig-3::HisCl1::SL2::mCherry];*<br>*[Pnpr-4-SNB-1::pHluorin; \*Lin-15]* | 6A, right panel |
| **AVA optogenetic (GtACR2) inactivation** |  |  |  |  |
| ZM10806 | *hpls824* | pJH4562<br>pJH4237 | *[Pflp-18-LoxP EBFP stop LoxP gtACR2::RFP;*<br>*Ptwk-40s-Cre]* | 3D; 3S4B |
| ZM10538 | *hpls774*;<br>*hpEx4081* | pJH4293<br>pJH4217<br>pJH4237 | <u>with AVA/AVB co-calcium imaging</u><br>*[Ptwk-40s-GCaMP6s::mNeptune]*<br>*[Prig-3 LoxP EBFP Stop LoxP gtACR2::RFP;*<br>*Ptwk-40s-Cre]* | 4B, 4S2B |
| **AVB optogenetic (GtACR2) inactivation** |  |  |  |  |
| ZM10942 | *lin-15(n765)*;<br>*hpls774*;<br>*hpEx4292* | pJH4293<br>pJH4609<br>pJH4237 | <u>with AVA/AVB co-calcium imaging</u><br>*[Ptwk-40s-GCaMP6s::mNeptune];*<br>*[Ppdf-1 LoxP eBFP LoxP gtACR2::wCherry;*<br>*Ptwk-40s-Cre; \*Lin-15(+)]* | 4A, B<br>4S2A |
| **AVB ablation (AVB miniSOG)** |  |  |  |  |
| ZM7297 | *hpls331* | pJH2890 | *[Plgc-55-mito::miniSOG UrSL wCherry;*<br>*\*Lin-15(+)]* | 4S1B |
| ZM10327 | *twk-40(bln282)*;<br>*hpls331*;<br>*hpls727* | pJH2890<br>pJH3830 | <u>with AVA-specific TWK-40(lf)</u><br>*twk-40::TagRFP::ZF;*<br>*[Plgc-55-mito::miniSOG sl2 RFP; \*Lin-15];*<br>*[Prig-3-zif-1 sl2 RFP]* | 4S1C |
| **AVA-AVB co-calcium imaging** |  |  |  |  |
| ZM10767 | *hpls819* | pJH4542 | *[Ptwk-40s-GCaMP T2A nls::mNeptune;*<br>*\*LIN-15(+)]* | 5A, B, D, E; 5S |
| ZM11014 | *twk-40(bln282)*;<br>*hpls819*;<br>*hpEx3904* | pJH4542<br>pJH4018<br>pJH2673 | <u>with AVA-specific TWK-40(lf)</u><br>*twk-40::TagRFP::ZF;*<br>*[Ptwk-40s-GCaMP:T2A:nls:RFP; \*Lin-15];*<br>*[Prig-3 sl2 FRT EBFP FRT zif-1;*<br>*Pnmr-1-FLP; \*HygR]* | 5C-E; 5S |
| ZM11015 | *hpls819*;<br>*hpls768* | pJH4542<br>pJH4316<br>pJH4237 | <u>with AVA-specific TWK-40(gf)</u><br>*[Ptwk-40s-GCaMP:T2A:nls:RFP; \*Lin-15];*<br>*[Pflp-18LoxPEBFPstopLoxP twk-40(L159N)::GFP*<br>*Ptwk-40s-Cre]* | 5F; 5SA, C |
| ZM11034 | *hpls819*;<br>*hpls810* | pJH4542<br>pJH4508<br>pJH4237 | <u>AVA-specific TeTx</u><br>*[Ptwk-40s-GCaMP:T2A:nls:RFP; \*Lin-15];*<br>*[Pflp-18-LoxP EBFP Stop LoxP TeTx::RFP;*<br>*Ptwk-40s-Cre]* | 6C |
| Notes: [ ] integrated (Is) or extra-transgenic (Ex) arrays; * Co-injection markers; Stop: stop codon or termination sequence in the let-858 3' UTR. |  |  |  |  |

---

**Figure 3S1.**
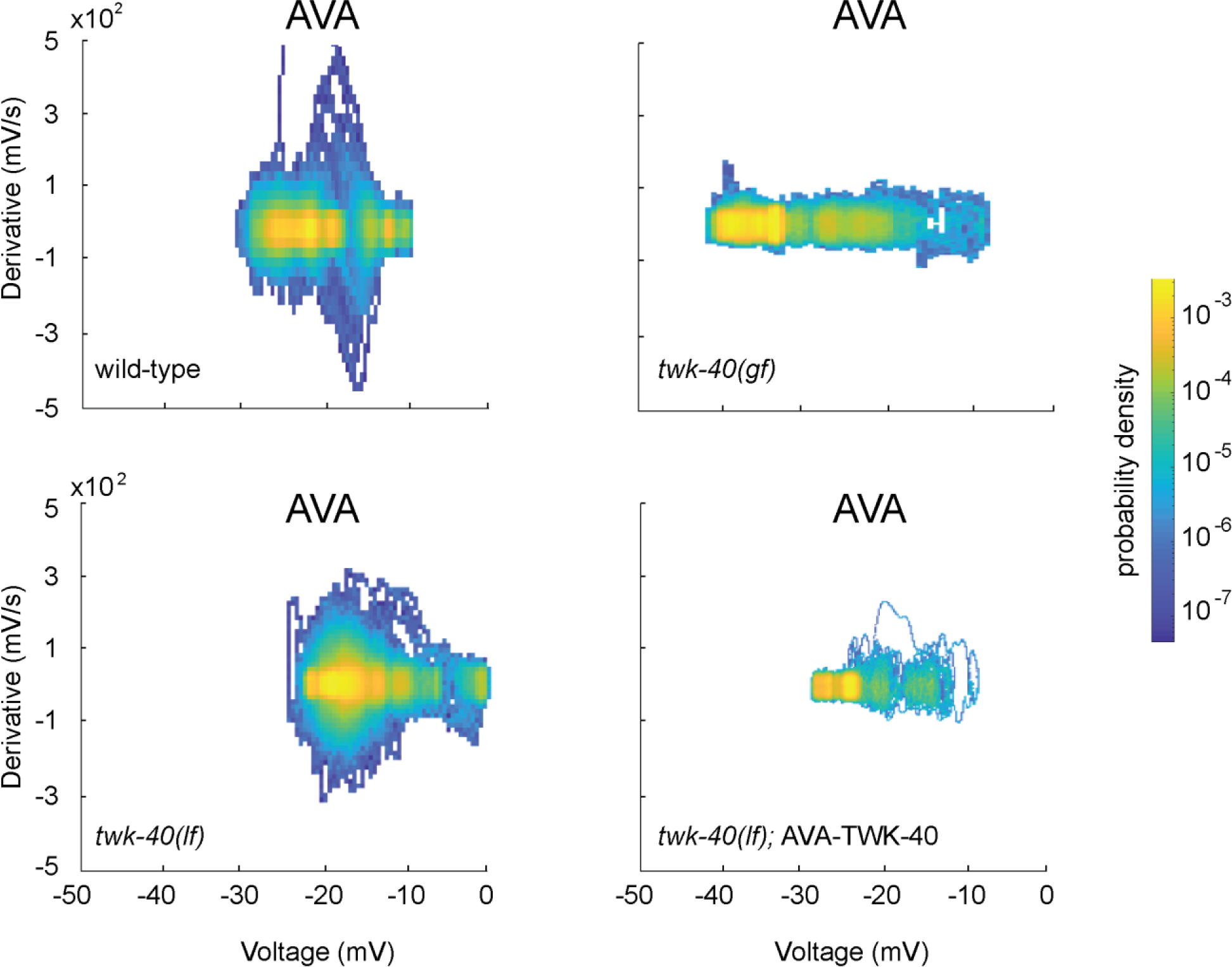
K+ leak regulates AVA’s membrane and dynamic range. Distribution of the voltage derivatives across AVA’s membrane potentials shows an increase and decrease of both the range and dynamic range in *twk-40(lf)* (upper right panel, N: 9 cells) and *twk-40(gf)* (lower left panel, N:13 cells) mutant animals, respectively. AVA-specific restoration of wild-type TWK-40 protein in *twk-40(lf)* mutants (lower right panel, N: 8 cells) decreased AVA’s range membrane potential and reduced its dynamic range, the latter similar to *twk-40(gf)* mutant animals. For each AVA neuron, recorded traces lasted for 60-200 seconds.

**Figure 3S2.**
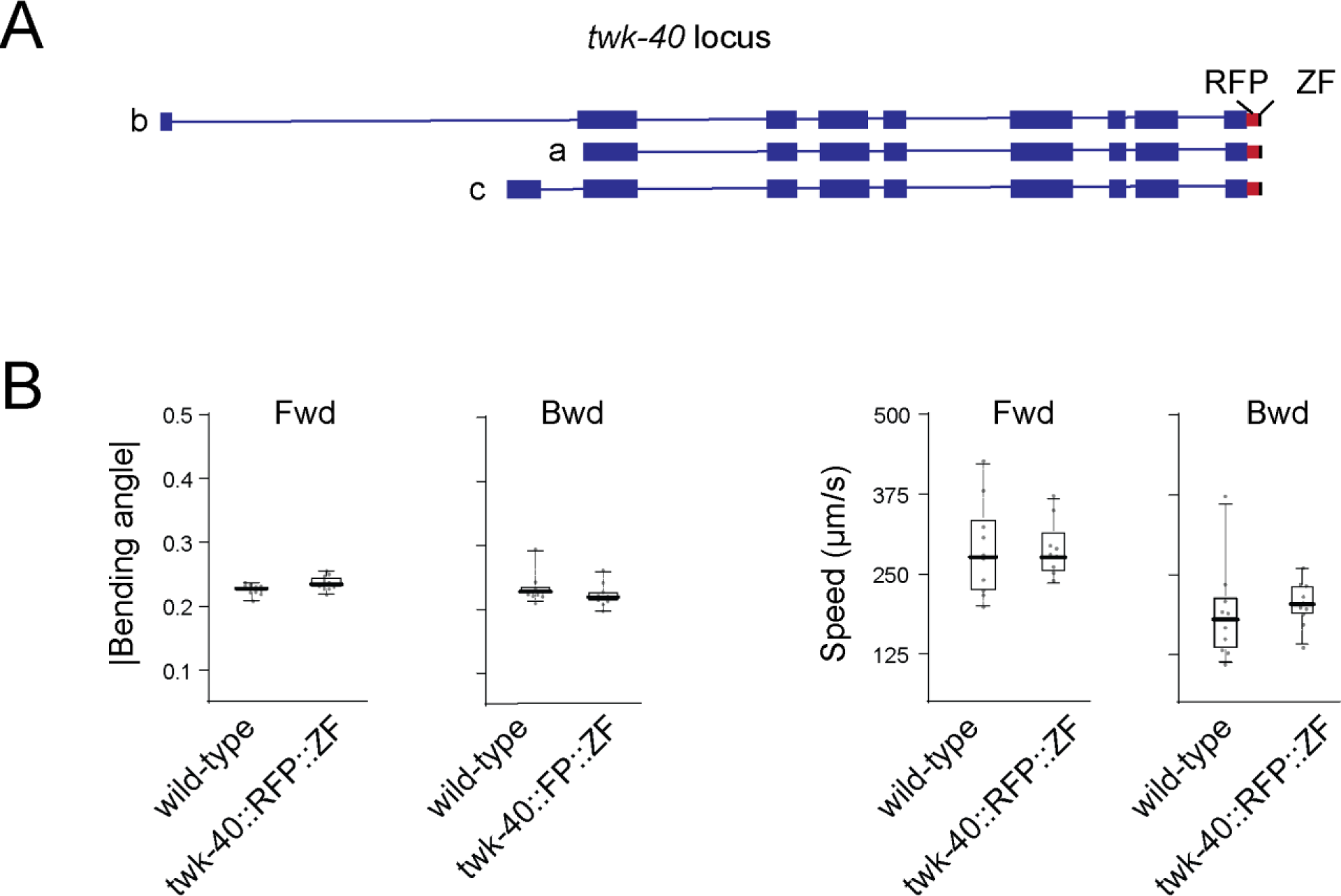
TWK-40::RFP::ZF knockin animals exhibit wildtype motor behaviors. **A.** Genomics structures of the *twk-40* gene, with three predicted protein isoforms (a, b, and c). Blue boxes and lines denote exons and introns, respectively. Red and green boxes denote an C-terminal in-frame insertion of the TagRFP and ZF domains in the endogenous *twk-40* genomic locus. **B.** Mean absolute bending angles (left) and centroid speed (right) of wild-type animals (N: 11) and *twk-40::TagRFP::ZF* animals (N: 11). Box-whisker plot displays the 90/10 and 75/25 percentiles, respectively. *: P < 0.05, **: P < 0.01, ***: P <0.001 by the Mann-Whitney U tests. P < 0.05 is considered to be statistically significant.

**Figure 3S3.**
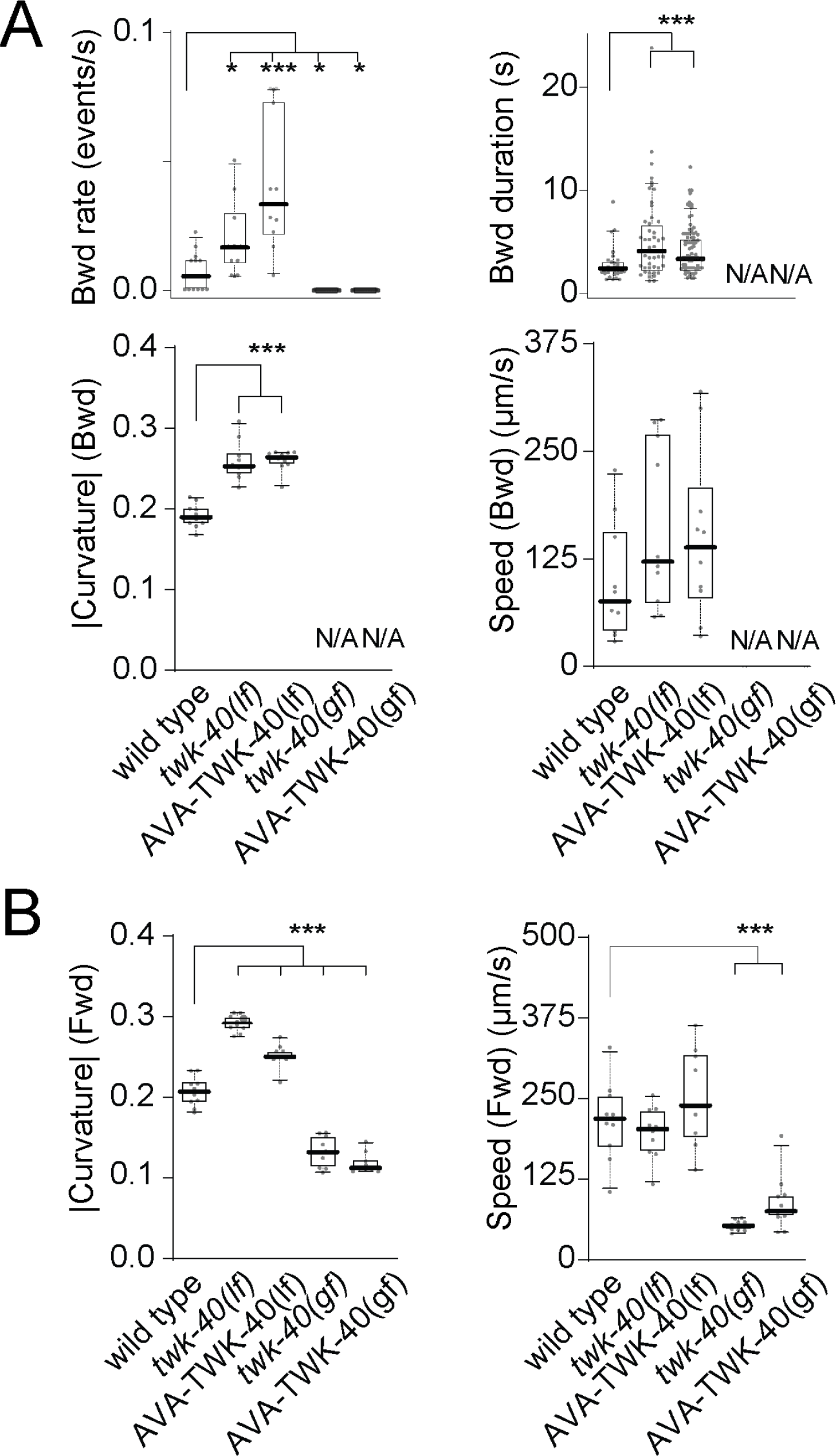
TWK-40 functions mainly through AVA to regulate locomotion. **(A)** Deceasing and increasing TWK-40 potassium leak led to increased and decreased backward movement. Quantification of (top left) backward movement rate, (top right) duration during backward movement, exhibited by animals of denoted genotypes. *twk-40(lf)* (N: 46 events from 11 animals) and AVA-specific TWK-40(lf) (N: 80 events from 11 animals) animals exhibit increased Backward movement rates and duration compared to wild-type animals (N: 30 events from 13 animals). (Lower left) Mean absolute bending angles and (lower right) centroid speed during backward movement. N: 11 (wild type), 11 (*twk-40(lf)*), 11 (AVA-specific TWK-40(lf)), 11 (*twk-40(gf)*) and 12 (AVA-specific TWK- 40(gf)) animals. **(B)** (Left) mean absolute bending angles and (right) centroid speed during forward movement by animals of denoted genotypes. N: 11 (wild-type), 11 (*twk-40(lf)*), 11 (AVA-specific TWK-40(lf)), 11 (*twk-40(gf)*) and 12 (AVA-specific TWK-40(gf)) animals. N/A: no events. Box-whisker plot displays the 90/10 and the 75/25 percentiles, respectively. *: P < 0.05, **: P < 0.01, ***: P <0.001 by the Mann-Whitney U tests. P<0.05 is considered to be statistically significant.

**Figure 3S4.**
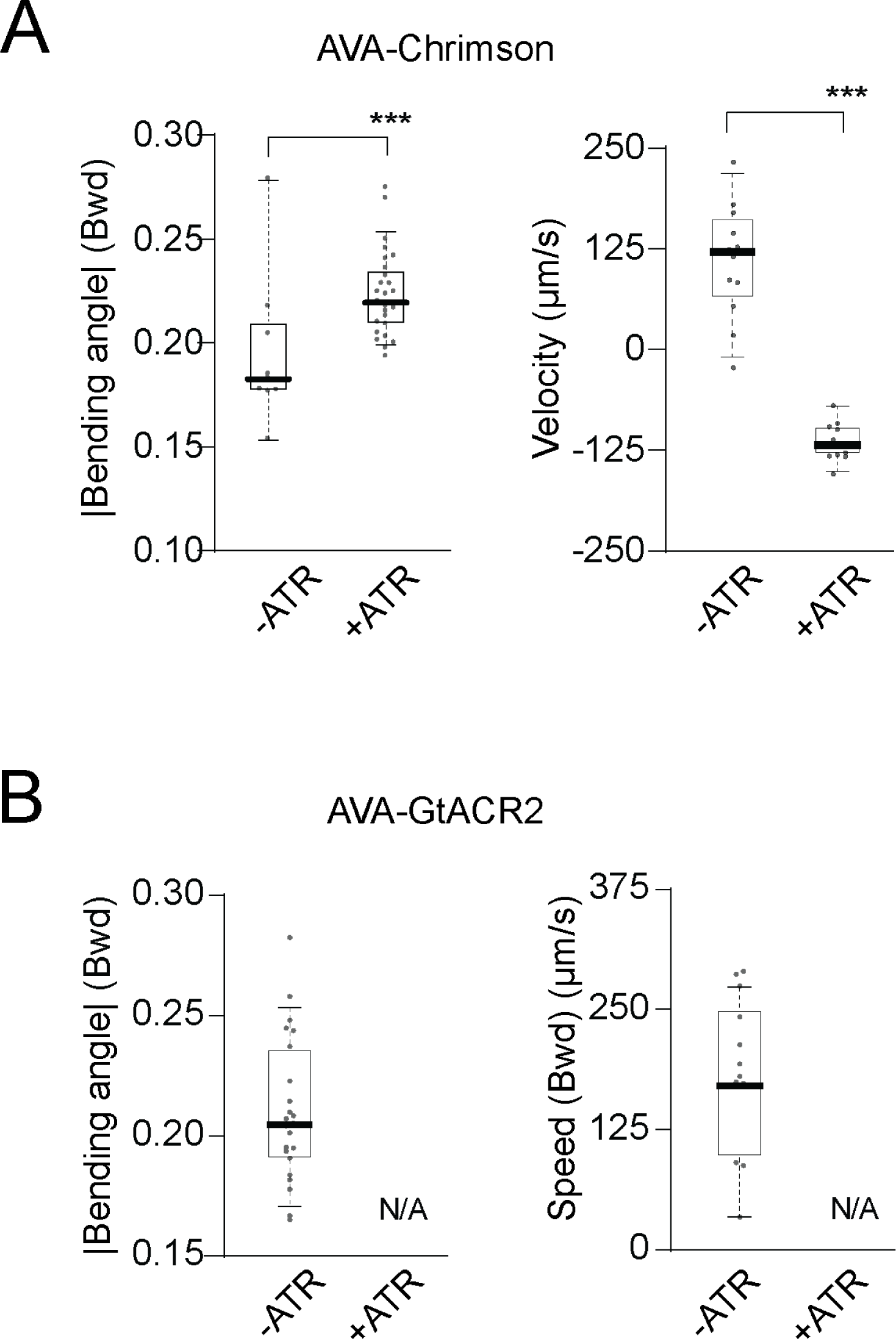
Optogenetic manipulation of AVA regulates the backward movement. **A.** Quantification of the backward movement, by the mean absolute bending angles (left) and centroid velocity (right), in non-stimulated (-ATR) (N: 28) and stimulated (+ATR) (N: 25) animals that express activating opsin (Chrimson) specifically in AVA. Acute AVA depolarization potentiates instantaneous backward movement. **B.** Quantification of the backward movement, by the mean absolute bending angles (left) and centroid speed (right), in non-stimulated (-ATR) (N: 12) and stimulated (+ATR) (N: 10) animals that express inhibitory opsin GtACR2 specifically in AVA. Backward movement was immediately and acutely blocked during stimulation. N/A: no events. **ATR**: all-trans retinal, co-factor for opsins expressed by transgenic *C. elegans*. Box-whisker plot displays the 90/10 and the 75/25 percentiles respectively. *: P < 0.05, **: P < 0.01, ***: P <0.001 by the Mann-Whitney U tests. P <0.05 is considered to be statistically significant.

**Figure 4S1.**
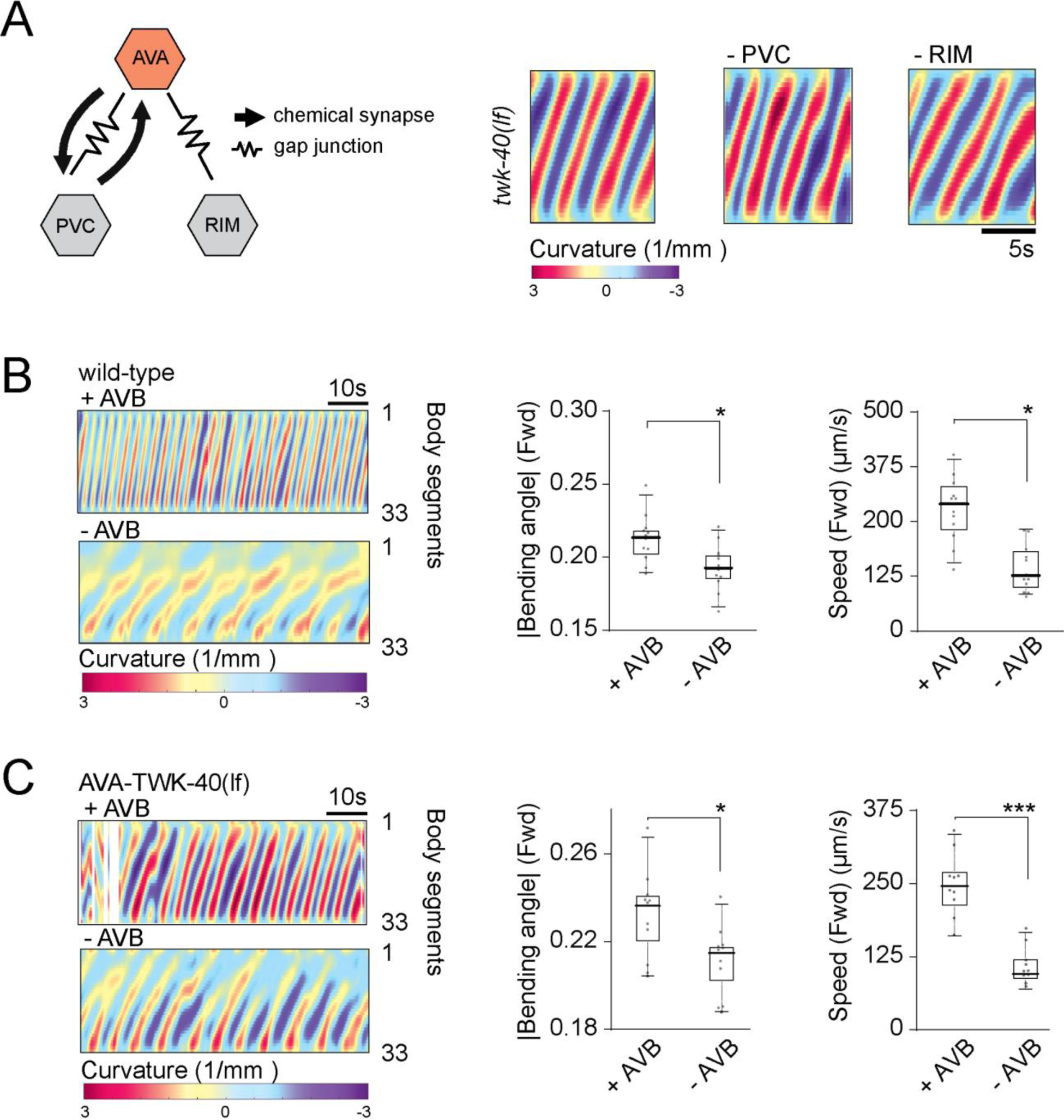
AVA-mediated potentiation of the forward movement requires AVB. **A.** (Left) Anatomic wiring diagram of other neurons that make consistent chemical and gap junctional connections with AVA, adapted from(*24*, *59*, *68*). (Right) Representative kymograph of *twk-40(lf)* mutant animal’s bending angles during forward movement, upon anatomic ablation of neurons of PVC or RIM, There were no significant changes in the bending angle or centroid speed. **B.** Anatomic ablation of AVB in wild-type animals led to significantly reduced forward bending angles and centroid forward speed. (Left) Representative kymographs of the bending angles of wildtype (control, +AVB; N: 13) and AVB ablated (-AVB) animals (N: 13). (Right) Mean absolute bending angles and centroid speed during the forward movement by animals of respective genotypes. **C.** Ablation of AVB led to significantly reduced bending angles and centroid speed during forward movement of AVA-specific TWK-40(lf) animals. (Left) Representative kymographs of bending angles of Control (+AVB, N: 11) and AVB ablated (-AVB) animals (N: 11) in AVA-specific TWK-40(lf) mutant animals. (Right) Mean absolute bending angles and centroid speed during the forward movement by animals of respective genotypes. Box-whisker plot displays the 90/10 and 75/25 percentiles, respectively. *: P < 0.05, **: P < 0.01, ***: P <0.001 by the Mann-Whitney U tests. P<0.05 is considered to be statistically significant.

**Figure 4S2.**
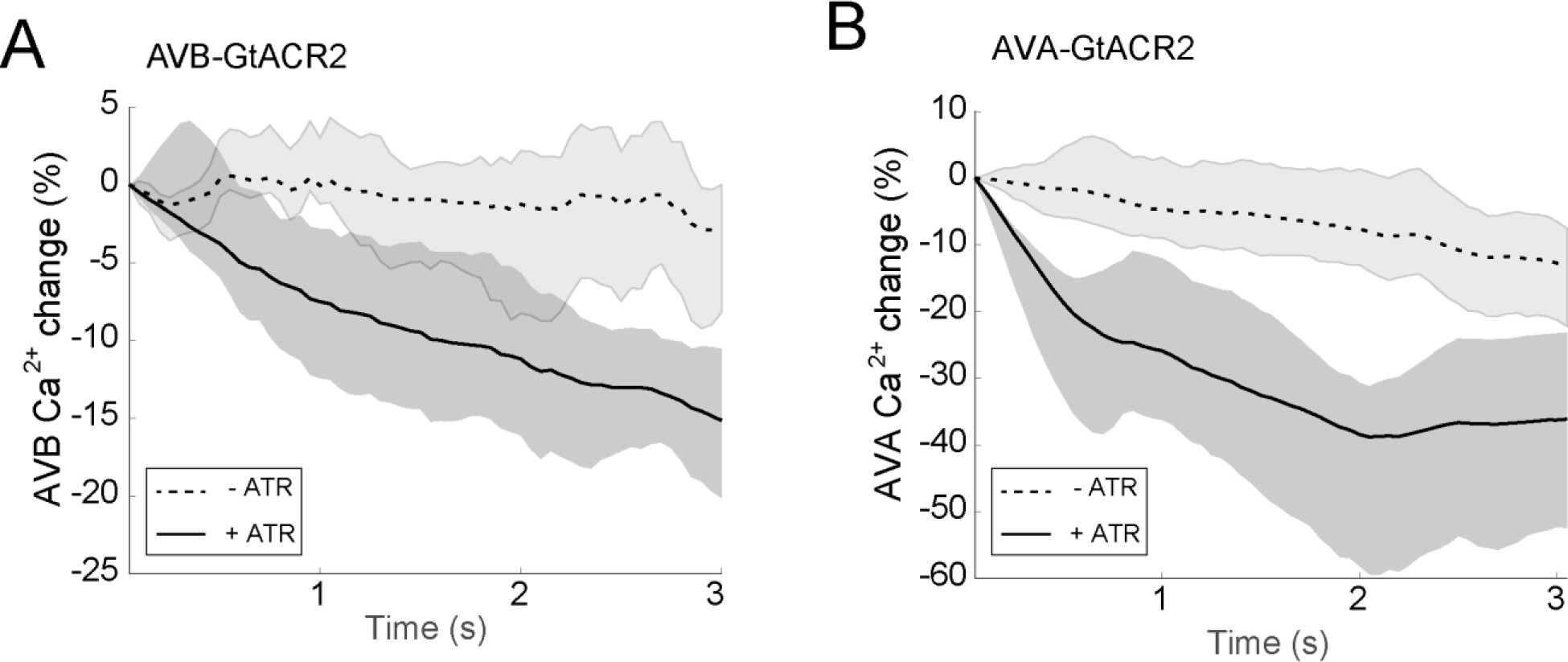
AVB-specific GtACR2 and AVA-specific GtACR2 are functional. **A.** Optogenetic inhibition of AVB by AVB-specific GtACR2 in immobilized animals led to reduced AVB calcium activity (+ATR; N: 28) compared to un-stimulated (-ATR; N: 19) animals. Lines and shaded bars denote the median and 95% confidence internal, respectively. **B.** Optogenetic inhibition of AVA by AVA-specific GtACR2 in immobilized animals led to reduced AVA calcium activity (+ATR; N: 23) compared to un-stimulated (- ATR; N: 23) animals. **ATR**: all-trans retinal, co-factor for opsins expressed by transgenic *C. elegans*. Lines and shaded bars denote the median and 95% confidence interval, respectively.

**Figure 5S.**
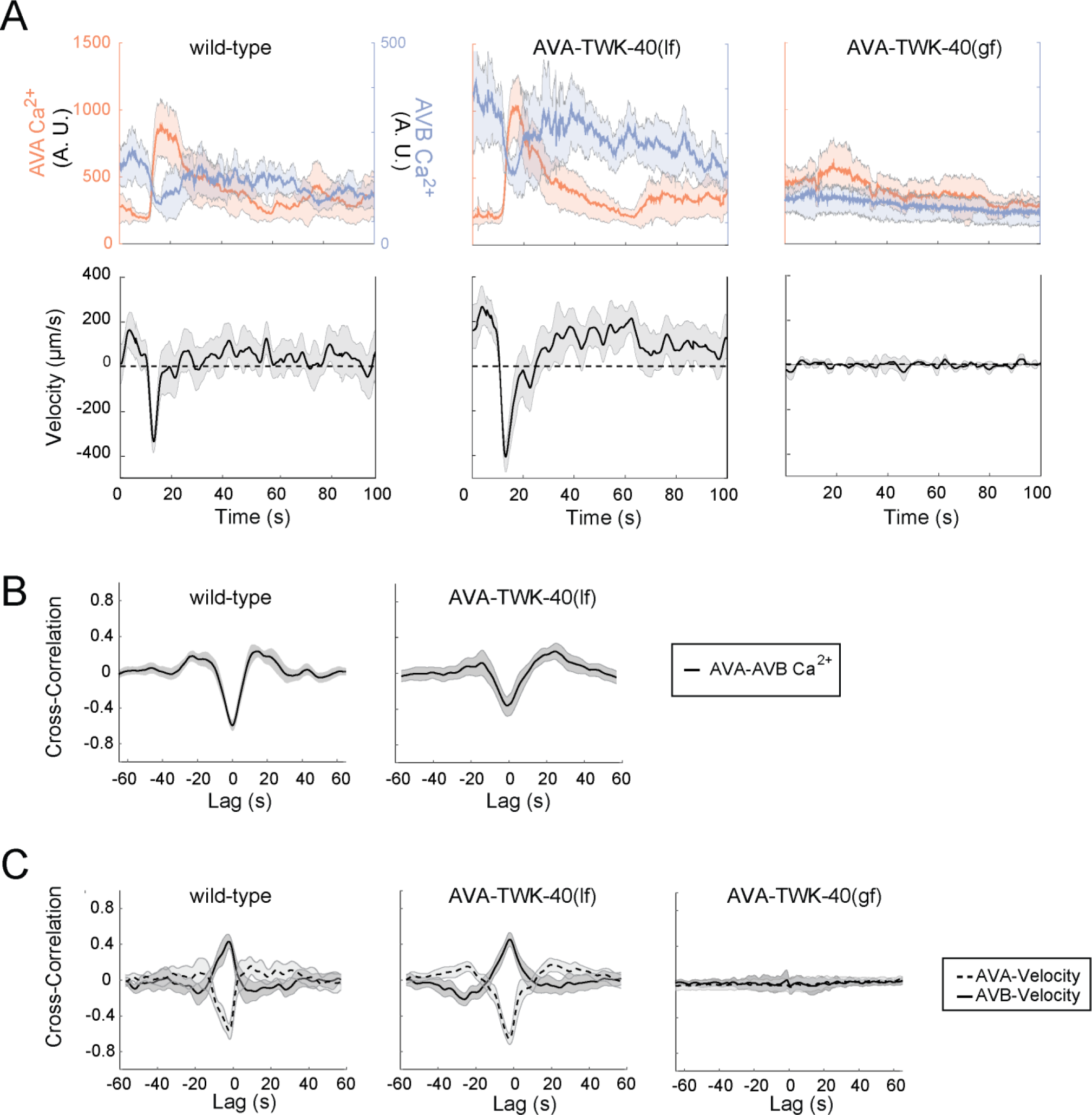
AVA’s activity is required to main the forward movement. **A.** (Upper panels) Mean calcium activity of AVA and AVB, aligned to the start of an AVA transient for wild-type, AVA-specific TWK-40(lf), and AVA-specific TWK-40(gf) animals, respectively. (Lower panels) Centroid velocities corresponding to the upper traces. + and – denote forward and backward movements, respectively. N: wild-type (19 epochs, from 11 animals); AVA-specific TWK-40 (27 epochs from 10 animals); AVA-specific TWK-40(gf) (4 epochs from 4 animals). Lines and shaded bars indicate the mean and 95% confidence interval. **B.** Mean cross-correlations between AVA and AVB’s calcium activity for wild-type (left panel) and AVA-specific TWK-40(lf) mutant (right panel) animals. **C.** Mean cross-correlations between AVA calcium activity and centroid velocity, and AVB calcium activity and centroid velocity for wild-type animals (Left panel), AVA-specific TWK-40(lf) mutants (Middle panel), and AVA-specific TWK-40(gf) mutants (Right panel). In both wild-type and AVA-specific TWK-40(lf) mutants, AVA calcium activity exhibited a negative correlation with backward speed during backward movement, and a weaker positive correlation with forward speed post-backward movement.

**Figure 6S1.**
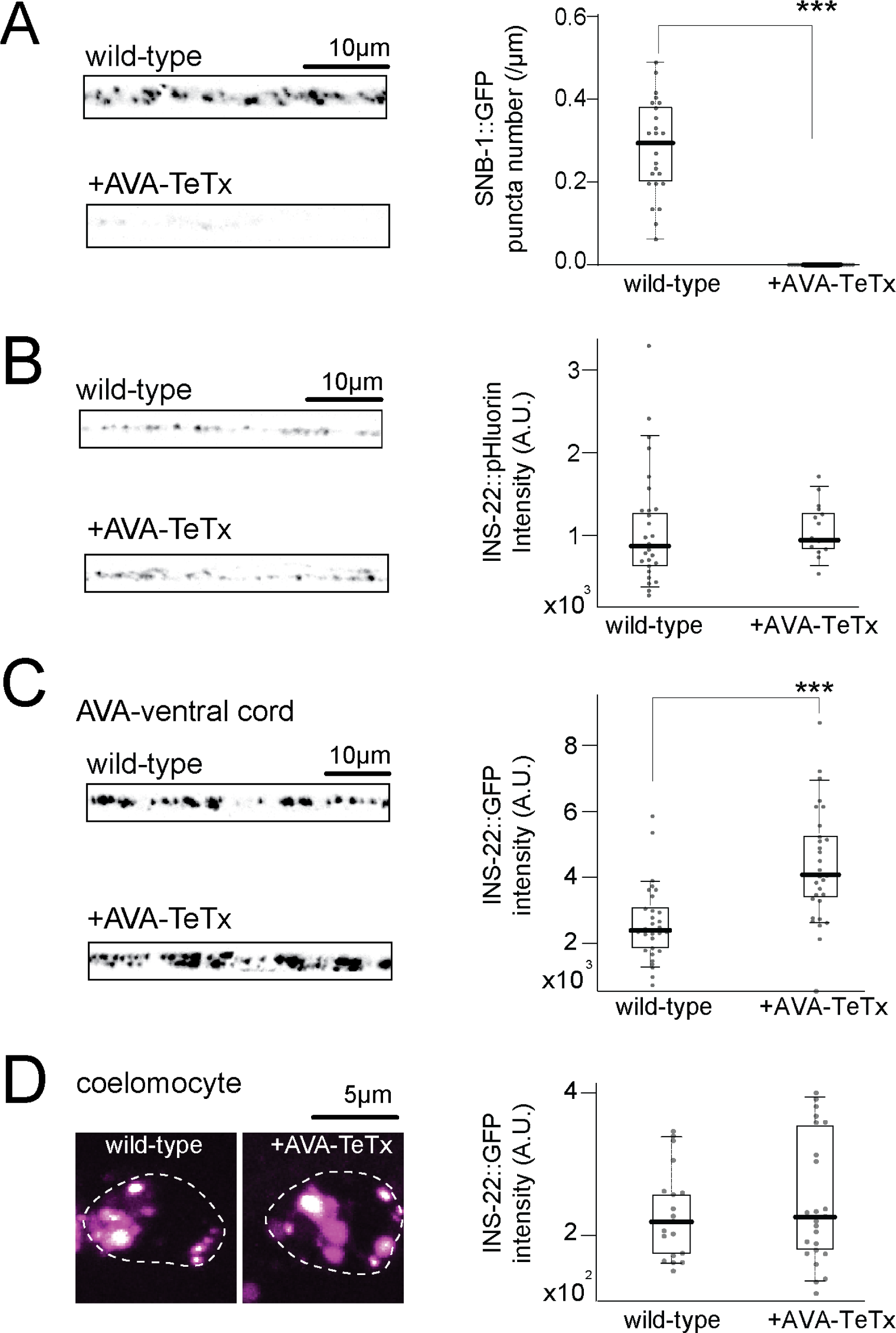
Tetanus toxin (TeTx), commonly used to block chemical synaptic transmission(*108*, *109*), cleaves synaptobrevin and preferentially blocks AVA’s chemical synaptic transmission. **A.** In AVA, TeTx potently cleaves SNB-1. Representative confocal images (Left panels) and quantifications of AVA-specific SNB-1::GFP signals (Right panel) in animals that express AVA-specific TeTx. SNB-1::GFP signals were diminished by the presence of TeTx. N: 22 AVA-TeTx animals and 25 wild-type animals. **B.** TeTx has a minor effect on neuropeptide secretion from AVA. Representative confocal images (Left panels) and quantification of fluorescent signals (Right panel) from an ectopically expressed INS- 22::pHluorin from AVA showed no significant change in the presence of TeTx. N: 15 AVA-TeTx animals and 27 wild-type animals. **C.** Representative confocal images (Left panels) and quantification of fluorescent signals of AVA-specific neuropeptide INS- 22::GFP marker (Right) showed increased intensity in the presence of AVA-specific TeTx. N: 30 AVA-TeTx animals and 30 wild-type animals. **D.** Representative images (Left panels) and quantification of secreted INS-22::GFP signals (Right), which accumulated in the coelomocytes. Mean INS-22::GFP intensity exhibited a wider range in AVA-TeTx animals (N: 26) but did not lead to a statistically significant difference with that of the wild-type animals (N: 20). Box-whisker plot displays the 90/10 and 75/25 percentiles, respectively. *: P < 0.05, **: P < 0.01, ***: P <0.001 by the Mann-Whitney U tests. P<0.05 is considered to be statistically significant.

**Figure 6S2.**
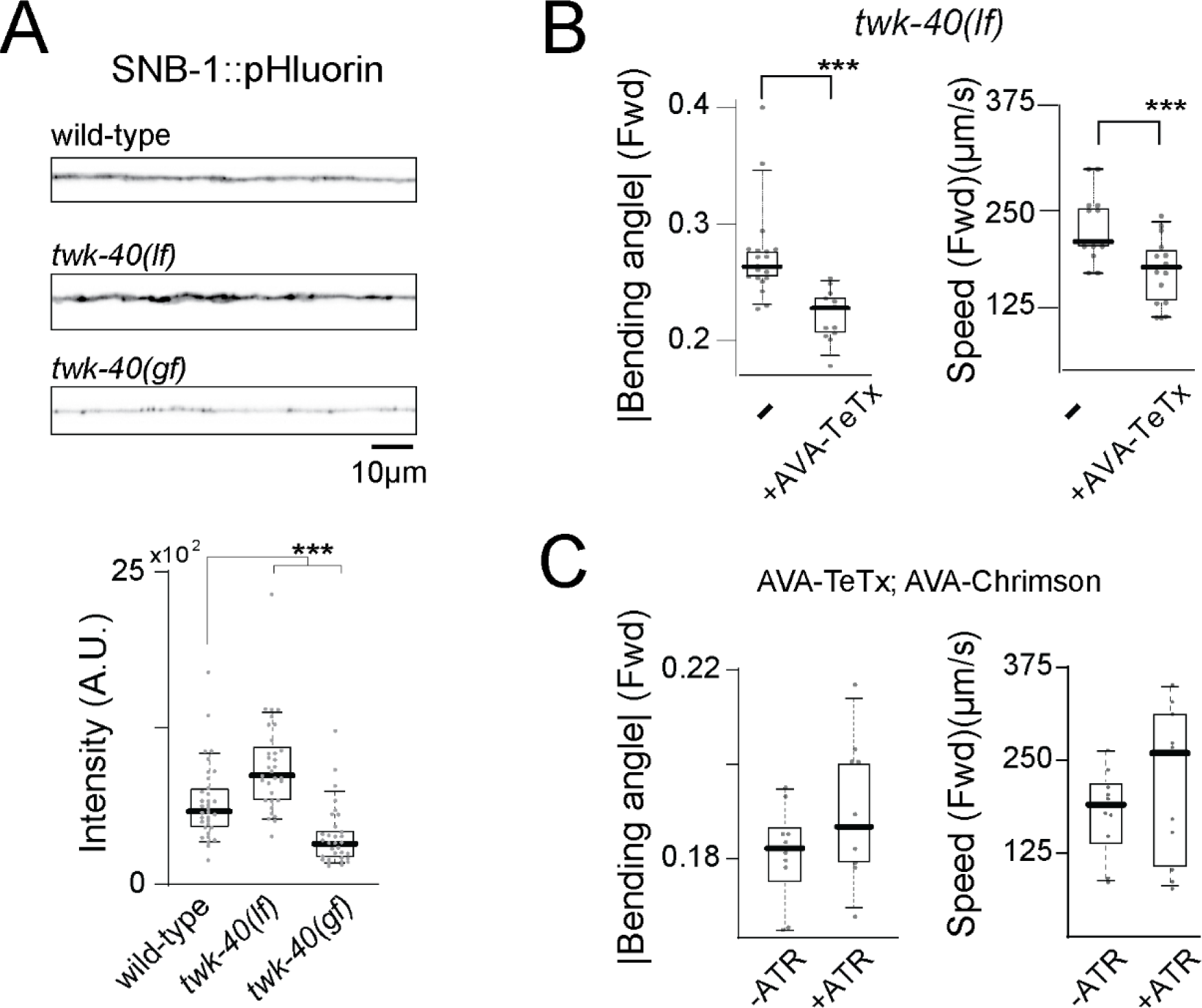
Acetylcholine release from AVA maintains the forward movement. **A.** Representative confocal images (Upper panels) and quantification (lower panel) of AVA-specific SNB-1::pHluorin signals in wild-type animals (N: 34), *twk-40(lf)* mutants (N: 32), and *twk-40(gf)* mutants (N: 32). pHluorin signals were significantly increased and decreased in *twk-40(lf)* and *twk-40(gf)*, respectively, compared to wild-type animals. **B.** Mean absolute bending angles (left panel) and centroid speed (right panel) during the forward movement by animals with synaptic vesicle fusion blocked (AVA-TeTx) in *twk-40(lf)* animals. The forward movement was reduced by AVA-specific TeTx. N: 20 *twk-40(lf)* and 14 AVA-TeTx; *twk-40(lf)* animals. **C.** Mean absolute bending angles (left panel) and centroid speed (right panel) of the forward movement upon AVA’s constitutive optogenetic activation (+ATR), in the presence of TeTx. Stimulation. They were compared with animals of the same genotype, treated with the same treatment but without opsin-activation (-ATR). When synaptic vesicle fusion was blocked, constitutive stimulation of AVA did not potentiate forward movement. N: 10 (-ATR) and 11 (+ATR) animals. Box-whisker plot displays the 90/10 and the 75/25 percentiles, respectively. **ATR**: all-trans retinal, co-factor for opsins expressed by transgenic *C. elegans*. *: P < 0.05, **: P < 0.01, ***: P <0.001 by the Mann-Whitney U tests. P<0.05 is considered to be statistically significant.

**Movie 1: Genetic manipulation of TWK-40 activity in AVA leads to robust changes in overall locomotory activity.** Representative movies of freely moving N2 (wild-type), *twk-40(lf)*, AVA-TWK-40(lf), *twk-40(gf)*, and AVA-TWK-40(gf) animals. AVA-specific degradation of TWK-40 leads to similar phenotype as *twk-40(lf)* genetic mutants, with increased body bending and velocity during both forward and backward movement. AVA-specific-expression of TWK-40(gf) leads to similar locomotor phenotype as *twk-40(gf)* mutant animals, with overall attenuation of body bending and velocity.

**Movie 2: AVA-specific optogenetic activation can mimic the locomotory phenotype of AVA-TWK-40(lf) animals.** Representative movies of freely moving animals expressing AVA-specific Chrimson, without and with ATR, the cofactor required for Chrimson activation. Acute and phasic Chrimson activation leads to instantaneous reversal, as reported previously. But with a moderate and constitutive activation, a locomotor phenotype similar to that of AVA-TWK-40(lf) animals, with an overall increase of body bending during forward and backward movement, was observed.

**Movie 3: AVA-specific optogenetic inactivation halts forward locomotion.** Representative movies of freely moving animals expressing AVA-specific GtACR2, without and with ATR, the cofactor required to activate GtACR2. Activation of GtACR2 led to instantaneous attenuation of the ongoing forward movement.

**Movie 4: Ablation of AVB reduces bending and velocity of spontaneous movement exhibited by *twk-40(lf)* and AVA-specific TWK-40(lf) animals.** Representative movies of freely moving animals with or without the ablation of AVB neurons in *twk-40(lf)* and AVA-TWK-40(lf) animals. ‘Non-ablated’ refers to animals that carried the miniSOG transgene for AVB ablation, but without carrying out the ablation. Compared to non-ablated controls, animals without AVB exhibited reduced bending and velocity.

**Movie 5. Ablation of PVC or RIM does not alter bending, velocity, or form of transitions during spontaneous movement in wildtype or *twk-40(lf)* animals.** Representative movies of freely moving animals with or without the ablation of either PVC or RIM neurons. ‘Non-ablated’ refers to animals carrying the respective miniSOG transgenes for PVC or RIM ablation, but without the ablation. ‘Wildtype’ refers to animals carrying the PVC or RIM miniSOG transgene. Ablation of PVC or RIM led to increased reversal frequency, but did not change the bending, velocity, or disrupt the smooth transitions between the forward and backward motor states, in both wildtype and *twk-40(lf)* animals.

## Methods

### Constructs and strains

A full list of constructs and stains used in this study is shown in Table 2. Constructs and strains are available at the Addagene, the *CGC* center, or upon request. All *C. elegans* strains were cultured on the standard Nematode Growth Medium (NGM) plates seeded with OP50 and maintained at 22.5 °C unless specified. The wild-type animal refers to the Bristol N2 strain or control animals as specified in figure legends. L4 stage hermaphrodites were used for all experiments unless specified otherwise.

### Isolation, cloning, and generation of *twk-40* mutants and AVA-specific TWK-40 alleles

The first *twk-40* loss-of-function(lf) allele, *hp733*, was isolated in a genetic screen based on altered motor behavior. *hp733* was isolated with exaggerated amplitude in both forward and reverse movement as well as a tendency to form a backward coil. *hp733* was mapped to the left arm of chromosome III past the SNP marker W06F12 at∼-21. Whole genome sequencing revealed a single missense mutation TWK-40(E36K) in *hp733* animals. Fosmids and plasmids containing the wild-type *twk-40* coding sequence fully rescued the motor phenotypes of *hp733*. We subsequently generated another *twk-40* lf allele, *hp834,* using CRISPR gene editing. *hp834* has a 7bp deletion in between exon 1 and the following intron (base 3378-3384 (GTTCGAG) of T28A8.1b), resulting in a frame-shift mutation of *twk-40* coding sequence that is predicted to translate only the first twelve amino acids of the channel. *twk-40(hp834)* mutant animals displayed motor behaviors of similar characteristics but more severe than that of *twk-40(hp733)* mutants. *hp834* was the canonical *lf* allele used for all experiments shown in this study.

Gain-of-function(gf) allele, *bln336(L181N)* was engineered by CRISPR/Cas9 editing by mutating the 4298-4299 base of T28A8.1b from CT to AA(*52*). Cell-specific TWK-40 degradation was generated by a repurposed ZIF E3 ligase degradation system(*55*). *bln282 (twk-40::TagRFP::ZF)* was generated by CRISPR/Cas9 genome editing to insert an C-terminal TWK-40 in frame TagRFP::ZF fusion at base pair 5785 at T28A8.1b(*104*). AVA-specific TWK-40(lf) animals were generated by expressing the ZIF E3 ligase specifically in AVA using the *Prig-3* or *Pnmr-1*/*Prig-3* combination (*Prig-3 FRT zif-1 sl2 eBFP FRT; Pnmr-1-FLP*) via the FRT-FLP recombination system in *bln282 (twk-40::Tag RFP::ZF)*.

AVA-specific TWK-40(gf) animals were generated by expressing TWK-40(L181N)::GFP specifically in AVA by P*flp-18*/P*twk-40* combination (*Pflp-18-LoxP stop LoxP twk-40(gf)::GFP; Ptwk-40-cre*) via the Cre-LoxP recombination system in the N2 background, or P*rig-3*/P*nmr-1* combination (*Prig-3-FRT stop FRT twk-40(gf)::GFP; Pnmr-1-FLP*) via the FRT-FLP recombination system or (*Pflp-18-LoxP stop Loxp twk-40(gf)::GFP; Ptwk-40-cre*) via the cre-Lox recombination system in N2 background. AVA-specific TWK-40 rescue was generated by expressing wildtype TWK-40(WT)::GFP specific in AVA by *Prig-3*/P*nmr-1* combination (*Prig-3 FRT Stop FRT twk-40::GFP; Pnmr-1-FLP*) via the FRT-FLP recombination system in *twk-40(lf)* background.

### Spontaneous locomotion analyses

A single L4 animal was collected and transferred to an NGM plate with a thin layer of evenly spread OP50. After 30 seconds, it was recorded on a tracking microscope using an in-house developed ImageJ plugin26. A 3-minute movie for each animal was recorded at 10 fps. At least 10 animals for each genotype were recorded on the same day using the same condition. Movies were analyzed as previously described(*67*, *71*). Briefly, the animal contour was divided into 33 segments by its centerline along the longitudinal axis. Body curvature in radian was calculated between adjacent segments for segments 5-32. Velocity was calculated by the displacement of the centroid.

### Electrophysiology

#### Exogenous TWK-40 channel whole-cell patch recording in HEK293 cells

HEK293 cells were cultured in the DMEM medium with 10% fetal bovine serum, 1% double antibody at 5% CO2, and 37°C. TWK-40 expression construct was generated by RT-PCR amplification of the *twk-40b* cDNA isoform and inserting into the BstXI and NheI sites of the pCDNA3.1 vector. After 24 hours of transfection of the plasmid with the liposome ExFect2000 (Vazyme, China), cells were patched using 4-6 MΩ-resistant borosilicate pipettes (1B100F-4, World Precision Instruments, USA). Pipettes were pulled by micropipette puller P-1000 (Sutter, USA), and fire-polished by microforge MF-830 (Narishige, Japan). Membrane currents and I-V curve were recorded and plotted in the whole-cell configuration by pulse software with the EPC-9 amplifier (HEKA, Germany) and processed with the Igor Pro (WaveMetrics, USA) and Clampfit 10 software (Axon Instruments, Molecular Devices, USA). I-V curve was recorded at a holding potential from –80 mV to 80 mV with 20 mV step voltages. Data were digitized at 10-20 kHz and filtered at 2.6 kHz. The pipette solution contained (in mM): KCl 140; MgCl2 1; EGTA 10; HEPES 10; Na2ATP 4; pH 7.3 with KOH, ∼300 mOsm. The bath solution consisted of (in mM): NaCl 140; KCl 3.6; CaCl2 2.5; MgCl2 1; pH 7.3 with NaOH, ∼310 mOsm. For low intracellular K+ recording, the pipette solution contained (in mM): KCl 14; CsCl 126; MgCl2 1; EGTA 10; HEPES 10; Na2ATP 4; pH 7.3 with CsOH, ∼300 mOsm. Chemicals were obtained from Sigma unless stated otherwise. Experiments were performed at room temperatures (20-22°C).

#### in situ whole-cell patch clamp of C. elegans neurons AVA and AVB

Premotor interneuron whole-cell patch clamp recording was performed using a strain with fluorescent labeling of AVA and AVB neurons, a configuration as previously described(*28*, *38*), which was modified from previous reports(*43*, *47*, *115*).

Briefly, 1- or 2-day-old hermaphrodite adults were glued (Histoacryl Blue, Braun) to a sylgard-coated cover glass covered with bath solution (Sylgard 184, Dowcorning) under a stereoscopic microscope (M50, Leica). After clearing the viscera by suction through a glass pipette, the cuticle flap was turned and gently glued down using WORMGLU (GluStitch Inc.) to expose the soma of AVA and AVB neurons. Neurons were patched using ∼20 MΩ-resistant borosilicate pipettes (1B100F-4; World Precision Instruments). Pipettes were pulled by micropipette puller P-1000 (Sutter) and fire-polished by microforge MF-830 (Narishige). Membrane currents and potentials were collected in the whole-cell configuration by pulse software with an EPC9 amplifier (HEKA, Germany). Data were digitized at 10 kHz and filtered at 2.6 kHz. The recording pipette solution consisted of: KCl 25 mM; K-Gluconate 115 mM; CaCl2 0.1 mM; MgCl2 5 mM; BAPTA 1 mM; HEPES 10 mM; Na2ATP 5 mM; Na2GTP 0.5 mM; cAMP 0.5 mM; cGMP 0.5 mM, pH 7.2 with KOH, ∼320 mOsm. cGMP and cAMP were included to increase the longevity of the preparation(*42*, *117*). The bath solution consisted of KCl 5 mM; NaCl 150 mM; CaCl2 5 mM; MgCl2 1 mM; glucose 10 mM; sucrose 5 mM; HEPES 15 mM, pH 7.3 with NaOH, ∼330 mOsm. The K+ current was recorded at a holding potential from –60 mV and followed a ramp voltage stimulation from –90 mV to 90 mV. Experiments were performed at room temperatures (20-22°C). Each recording lasted 60-200 seconds, with>90% at 200 seconds.

Membrane voltage recording was performed at 0 pA for at least 30 seconds before data collection. Healthy preparations were selected based on the following criteria: whole-cell capacitance (1-1.5 pF) and steady-state leak current (–20-0 pA at –60mV). As in our previous recordings, cAMP and cGMP were included in the pipette solution to maintain the activity and longevity of the preparation. Short traces (up to 200 seconds) without obvious rundown or re-sealing were used for analyses (*39*, *120*).

### Cell ablation by miniSOG

Strains expressing miniSOG(*116*) in AVB (and other neurons) or RIM (specifically) were used for ablation by exposing the respective L2 larvae to blue LED light as described in previous studies(*27*, *67*, *117*). Briefly, cells expressing miniSOG were co-labeled with RFP.animals were treated with 30-40 min blue light using a homemade LED device and allowed to recover for 1 day. L4 animals were recorded for behavioral analyses. For PVC ablation, L2 transgenic animals that express miniSOG in multiple premotor interneurons, including but not exclusive to PVC, were anesthetized by sodium azide and mounted onto a 2 percent agarose pad with a coverslip. The tail region, where PVC resides, was exposed to a blue laser for 30 min and animals were allowed to recover for 2 days before recording at the L4 stage. Only healthy animals were recorded and analyzed. The ablation efficiency and specificity (for PVC) for all experiments was confirmed by the disappearance or distortion of the soma fluorescence and fragmentation of neurites.

### Optogenetic stimulation for behavioral and calcium imaging

#### Strains

Strains that express AVA-specific Chrimson or GtACR2 strains were used for AVA-specific stimulation and inhibition, respectively. The specificity of AVA-specific GtACR2 was verified by AVA ablation using *Pnmr-1*-miniSOG. The AVA-ablated animal no longer exhibited light-induced inhibition during forward and backward movement.

Opsins expressed by transgenic *C. elegans* require exogenously supplied co-factor, all-trans retinal (ATR) for function. Transgenic animals were cultured on ATR plates in darkness for two generations. The second-generation L4-stage animals were analyzed for behavior and calcium activities. Controls were animals of the same genotypes, cultured and treated similarly, except with no ATR included on culture plates.

#### Behaviors upon stimulation

an individual L4-stage animal was transferred to an imaging plate with a thin layer of evenly spread OP50. LED light stimulation were applied to inhibit or activate AVA, by activating GtACR2 or Chrimson, respectively as below. *GtACR2-mediated AVA inactivation:* Animals were cultured on with or without ATR- supplemented NGM plates for 2 hours. During stimulation, L4 stage animals were exposed to 470nm LED light (Thorlabs) at low intensity (4 out of 63 A.U.) with a 10 second pre-stimulation, 10 seconds stimulation and 10 second post stimulation cycle. *GtACR2-mediated AVB inactivation:* L4 stage AVB-specific GtACR2 animals were incubated on plates with or without -ATR or +ATR (1ml OP50 /5μl 50μM ATR) plates for 2 hours and then subjected to behavior recording with a constant 470nm LED light (Thorlabs) exposure at low intensity (4 out of 63 A.U.) for 800 seconds.

#### Chrimson-mediated AVA activation

Animals were cultured on with or without ATR- supplemented NGM plates for at least 1 day for behavior recording. During stimulation, L4 stage animals were exposed to 574nm LED light (Thorlabs) at low intensity (63 out of 63 A.U.) with a 10 second pre-stimulation, 10 seconds stimulation and 10 second post stimulation cycle.

#### Constitutive, moderate Chrimson-mediated AVA activation

with or without ATR- supplemented NGM plates were seeded with regular OP50 supplemented with ATR (1ml OP50/0.5μl 50μm ATR). Animals were cultured on plates for at least 1 day for behavior recording. Before recording, (L4 stage) AVA were constitutively activated by 544nm green LED light for up to 5min (Zeiss V16), followed by recording at 595nm LED (Thorlabs) at low intensity (4 out of 63 A.U.).

#### Simultaneous calcium recording and optogenetic stimulation

The experiment process was previously described(*117*). Briefly, L4 stage animals carrying AVA and AVB calcium reporters and either AVA-specific GtACR2 or AVB-specific GtACR2 were incubated on - ATR or +ATR plates for 2 hours. They were immobilized on dry agarose pads in M9 buffer and subjected to full spectrum light illumination for simultaneous calcium imaging and GtACR2-mediated inhibition at 10 fps or 20 fps for 6-20 seconds.

### Behavioral analyses upon chemogenetic manipulation

Animals of the L4 stage carrying an integrated chromosomal array for *Prig-3* HisCl::mCherry(*36*) were incubated on 0mM, 2mM, 5mM, or 10mM histamine plates with OP50 for 1 hour before being individually placed on the behavioral imaging plates for spontaneous behavior recording and quantification.

### Immunofluorescent staining and imaging

Mixed stages of animals with endogenously tagged *twk-40* (*twk-40*::TagRFP::ZF), with or without the plasma marker (*Prgef*-1-GPI::YFP), and animals with an integrated *twk-40::GFP* fosmid were fixed in 2% paraformaldehyde at 4°C for 1 hour and processed as previously described(*118*). Primary antibodies against TagRFP (Thermo Fisher Scientific, R10367, rabbit) and/or GFP (Roche, mouse) were used at 1:500 and 1:200 dilution, respectively. Secondary antibodies of goat-anti-rabbit (Alexa Fluor 488) and/or goat anti-mouse (Alexa Fluor 594) were used at 1:5000 dilution. L4 larvae and adults were imaged with a Nikon spinning disk confocal microscope with a 63x objective and reconstructed by maximum intensity projection. Single layers of acquired images from animals co-expressing TWK-40::TagRFP::ZF and GPI::YFP were examined for subcellular colocalization using an ImageJ plugin (JACoP).

### Live fluorescence microscopy of GFP and pHluorin

Live L4 larvae were immobilized on dry agarose pads in a small drop of M9 buffer. Fluorescence signals were captured using Nikon Eclipse 90i confocal microscope and Nikon Eclipse T12 with Yokogawa CSU-X1 spinning disk field scanning confocal system. For pHluorin imaging, all images were acquired within 10min after dry pad preparation. All fluorescent images were analyzed by the in-house developed puncta analyzer (*119*) or ImageJ intensity analyzer.

### Calcium imaging of premotor interneurons in moving animals

For AVA (L or R) and AVB (L or R) co-imaging, an individual transgenic L3 stage animal expressing the GCaMP6s calcium sensor in AVA and AVB (*hpIs819*) was mounted on a 2 percent NGM gel pad with ∼1μl M9 buffer so that it could move under a coverslip.

Mounted animals were allowed to recover for 3min on the pad before recording. The recording was performed using a 20x objective on Nikon spinning disk confocal microscope. Animals were centered in the field of view by manual tracking. AVB soma was manually kept in focus at the RFP channel during recording. The initiation of each epoch of backward to forward locomotion transition was defined by the alignment between velocity and AVA-AVB calcium activity (GCaMP intensity) using in-house-developed MATLAB scripts(*117*).

### Cell-specific perturbation of neurotransmission

AVA-specific TeTx was achieved by the Cre-LoxP recombination system. AVA-specific *cha-1* RNAi was generated using the combination of P*flp-18* and *Ptwk-40* promoter combination to drive the sense and anti-sense strands of *cha-1* genomic DNA, respectively. The following primers and templates were used:

(A) *Pflp-18*: OZM6199 (GATGGATACGCTAACAACTTGG (OZM6199) and OZM6200 (CTACAACGGCAGCGTATTCGATCCCGTCTAACCCTGAAA) amplified from pJH4631.

(B) *cha-1* exon: OZM6201 (GAATACGCTGCCGTTGTAG) and OZM6240 (GTCGAGTGCTCTATGCACAACC), amplified from N2 genomic DNA.

(C) *Ptwk-40s*: OZM6199 (GATGGATACGCTAACAACTTGG) and OZM6253 (GTATGATGCGACTATTCAGCTGT-GAATATTCATCACTCGATATTCCA) amplified from pJH4608.

(D) *cha-1* exon: OZM6201 (GAATACGCTGCCGTTGTAG) and OZM6202 (CAGCTGAATAGTCGCATCATAC) amplified from N2 genomic DNA.

PCR fusion was performed to obtain sense and anti-sense *cha-1* by following primers*: Pflp-18-cha-1*(sense): OZM6203 (AGCTTAGCCGGAATAGGGTCA) and OZM6202 (CAGCTGAATAGTCGCATCATAC), amplified from mixed PCR product (A+B).

*Ptwk-40-cha-1*(antisense): OZM3543 (AATGCGGCCGCTCTAATCACTATCACGT GGGATCTGGATAA); OZM6254 (CGTAG-GCCAGAAAGCCTCAC) from mixed PCR product (C+D). Mixed PCR products were injected into N2 animals.

### Modeling

Our approach is to find the simplest phenomenological model (i.e. a non-mechanistic model with as few free parameters as possible) that explains the observed calcium activity patterns of AVA and AVB, as opposed to a detailed, more biologically realistic model. A more realistic model will have several parameters, the estimation of which will require assumptions or experiments to fix.

We first note that our experiments suggest that the communication between AVB and AVA is largely unidirectional, with AVA affecting AVB’s activity and very little effect in the opposite direction. Consequently, we start with the following equation for the calcium activity of AVB, *x_AVB_*

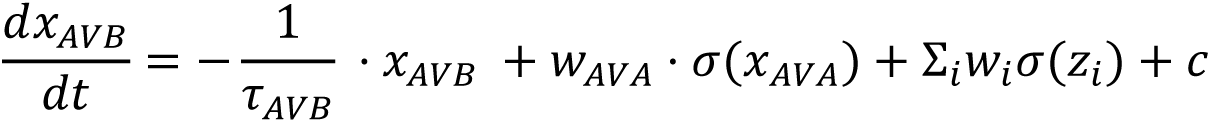

Where *x_AVA_* is the calcium activity of AVA, *z_i_* is the activity of all other input neurons of AVB, *w_i_* is the synaptic weight of each input, *c* is the bias input, and *σ* denotes the activation function for the synapse. Typically, the activation function is a sigmoidal function. However, to simplify the model, we restrict our attention to the linear case, where *σ* is simply the identity function, *σ(x) = x*.

Next, we make a key simplification to the model above based on our experiments. Our results show that AVA forms the dominant input on to AVB. Specifically, AVA-twk-40(gf) animals cannot sustain their forward motion (Fig. 2B) and AVB’s calcium is significantly reduced in this background (Fig. 3D). This suggests that we can ignore the input from other neurons in the model. Thus, we set *w_i_* = 0 for all neurons other than AVA. Giving us the following leaky-integrator differential equation.

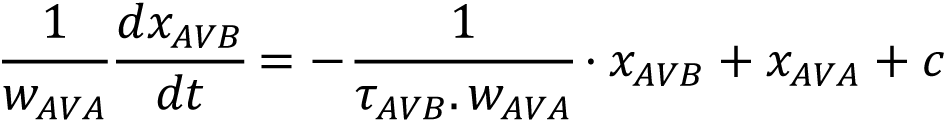

As a final simplification, we assume that the direct input from AVA is faster than the kinetics of the calcium sensor, such that any activity in AVA is immediately reflected in AVB. Thus, we consider the limit 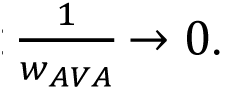 xGiving us the final simplified model as

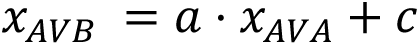

Where 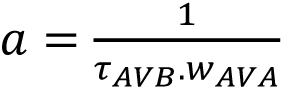 is a dimensionless multiplier. As seen in Fig. 5B, this model with *a* <0 (black dotted line) can predict the anticorrelated activity pattern between AVA and AVB during a backward movement, but is unable to explain AVB’s activity after the backward movement.

To capture the slow decay of AVB, we add an additional slower input from AVA to AVB (as shown in the schematic in Fig. 5A). Giving us the following system of equations with just three free parameters.

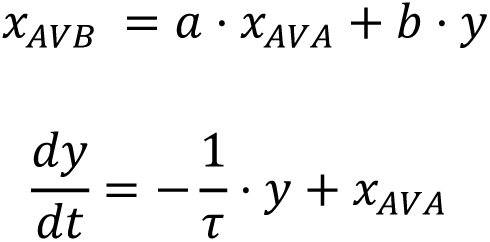

Where, *a* and *b* are the coefficients for the fast and slow inputs respectively, and *τ* is the integration time-constant for the slow component of AVA. To fit the parameters from calcium imaging data, we used *procest* function from MATLAB’s system identification toolbox, which estimates the transfer function for the equation above. Laplace transforming the system of equations above results in the following transfer function,

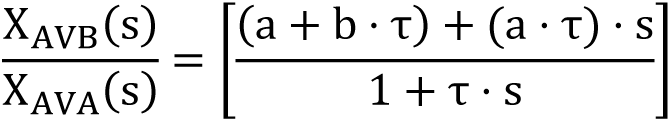

which has one zero and one pole. The number of zeros and poles form the input to *procest*, along with the calcium traces of AVB calcium (set as output signal) and AVA calcium (set as the input signal). To get a distribution over model parameters and performances, we do a cross-validation procedure where the parameters are estimated from a single recording and then tested on the remaining recordings. The model R-squared showed in Fig 5C shows the distribution from the models thus estimated. The same setup was used to model the velocity from AVA’s calcium activity (Fig 5E) using *procest* (i.e. a single pole, single zero transfer function), with AVA’s activity being set as the input signal and velocity forming the output signal.

### Statistical analyses

#### Electrophysiological analyses

The cumulative distributions for the membrane voltage of AVA and AVB were calculated using MATLAB’s *ecdf* function. To compute the joint histogram (Fig. 1S1, Fig. 3S2), we first computed the derivative of membrane potential with a finite difference (preceded by smoothing with a Gaussian kernel) and then used MATLAB’s bivariate histogram *histogram2* to perform the histogram estimation.

#### HEK cell recording, Behavioral analyses, and fluorescence staining

Mann-Whitney U tests were used to quantify data of box and whisker plots.

#### Calcium imaging

Error bars in Fig. 4 and Fig. 4S were computed as a 95% confidence interval of the mean bootstrapped over 1000 resamples.

P value less than 0.05 is considered to be statistically significant. *: P < 0.05, **: P < 0.01, ***: P <0.001.

## Author contributions

Conception and supervision: MZ; Design of experiments, modeling, and analyses: JM, TA, WH, SG, TB, MZ; Data collection: JM, BY, WH, YZ, SG; Reagents and scripts: JM, WH, SM, ZW, YZ, TA, QW; JM, TA and MZ wrote the manuscript; all authors edited or approved the manuscript.

## Data availability

All data needed to evaluate the conclusions in the paper are present in the paper and/or the Supplementary Materials.

## Acknowledgments

We thank Ying Wang and Yan Li for technical support; comments from and discussions with Aravithan Samuel, Ben Mulcahy, Miriam Goodman, and Cori Bargmann. This work was supported by the Canadian Institute of Health Research FDN154274 and the Natural Sciences and Engineering Research Council of Canada RGPIN2017-06738 (MZ), the key International (Regional) Joint Research Project (32020103007) (SG and QW), the National Natural Science Foundation of China (31871069) (SG) and the European Research Council (337702-Kelegans) (TB).

